# Enrichment of gene variants associated with treatable genetic disorders in psychiatric populations

**DOI:** 10.1101/287219

**Authors:** Venuja Sriretnakumar, Ricardo Harripaul, John B. Vincent, James L. Kennedy, Joyce So

**Affiliations:** Campbell Family Mental Health Research Institute, Centre for Addiction and Mental Health, 250 College Street, M5T 1R8, Toronto, Canada; Department of Laboratory Medicine and Pathobiology, University of Toronto, Medical Sciences Building, 6th Floor, 1 King’s College Circle, M5S 1A8, Toronto, Canada; Institute of Medical Science, University of Toronto, Medical Sciences Building, Room 2374, 1 King’s College Circle, M5S 1A8, Toronto, Canada; Department of Psychiatry, University of Toronto, 250 College Street, 8th floor, M5T 1R8, Toronto, Canada; The Fred A. Litwin Family Centre in Genetic Medicine, University Health Network and Mount Sinai Hospital, 60 Murray Street, M5T 3L9, Toronto, Canada; Department of Medicine, University of Toronto, Toronto, Canada

**Keywords:** inborn errors of metabolism, schizophrenia, bipolar disorder, major depressive disorder, psychiatric genetics

## Abstract

**Purpose:** Many genetic conditions can mimic mental health disorders, with psychiatric symptoms that are difficult to treat with standard psychotropic medications. This study tests the hypothesis that psychiatric populations are enriched for pathogenic variants associated with selected treatable genetic disorders.

**Methods:** Using next-generation sequencing, 2046 psychiatric patients were screened for variants in genes associated with four inborn errors of metabolism (IEMs), Niemann-Pick disease type C (NPC), Wilson disease (WD), homocystinuria (HOM), and acute intermittent porphyria (AIP).

**Results:** Among the 2046 cases, carrier rates of 0·83%, 0·98%, 0·20%, and 0·24% for NPC, WD, HOM, and AIP were seen respectively. An enrichment of known and likely pathogenic variants in the genes associated with NPC and AIP was found in the psychiatric cohort, and especially in schizophrenia patients.

**Conclusion:** The results of this study support that rare genetic disease variants, such as those associated with IEMs, may contribute to the pathogenesis of psychiatric disorders. IEMs should be considered as possible causative factors for psychiatric presentations, especially in psychotic disorders, such as schizophrenia, and in the context of poor treatment response.

## INTRODUCTION

Schizophrenia (SCZ) is a debilitating psychiatric disorder characterized by positive (e.g. hallucinations) and negative (e.g. avolition) symptoms, and cognitive deficits, with a global prevalence of up to 1·1%.^1^ One-third of SCZ patients are non-responsive to standard anti-psychotic medications.^1^ Bipolar disorder (BPD) is an affective disorder characterized by periods of mania and depression, with a lifetime prevalence of up to 4%.^2^ Up to 37% of BPD patients are treatment-resistant to first-line mood-stabilizing drugs.^2^ Major depressive disorder (MDD) is the most common psychiatric disorder worldwide with a 16·2% lifetime prevalence and poor treatment response in up to 30% of patients.^3^ SCZ, BPD, and MDD have high heritability rates upwards of 0·85, 0·79, and 0·45, respectively.^4^ Large concerted efforts to determine the causative genes for SCZ, BPD, and MDD through genome-wide association studies have identified numerous genes with only small effect sizes.^1-4^ These studies have focused on identifying common variants, with less attention being attributed to rare, more penetrant variants.

Though individually rare, genetic diseases are collectively common, with an estimated global prevalence of 1 in 100.^5^ Often overlooked is that many genetic diseases are treatable, particularly inborn errors of metabolism (IEMs), a group of rare diseases that result from mutations in genes that encode proteins or enzymes along metabolic pathways. Although most IEMs are diagnosed in early childhood, there has been an increased recognition of late- or adult-onset IEMs that can present with psychiatric symptoms that are indistinguishable from primary psychiatric illnesses, such as SCZ; examples of such disorders are shown in supplementary Table 1.^6^ Significantly, identification of a genetic diagnosis in a psychiatric patient would reveal targeted management and, particularly with IEMs, specific treatments that could ameliorate the psychiatric symptoms and prevent onset of other systemic features of the diseases. To date, there have been several literature reviews and case studies suggesting the importance of investigating IEMs as a primary factor in psychosis and SCZ (e.g. Nia *et al.* (2014)).^6^ Trakadis *et al.* (2017) recently reported a significant mild enrichment (odds ratios between 1·13-1·64) in an unselected group of 2545 adults with SCZ from the Database of Genotypes and Phenotypes of rare, presumed functional variants in several IEM genes, including *NPC1* and *NPC2* (associated with Niemann-Pick disease type C (NPC)), *ATP7B* (associated with Wilson disease (WD), and *CBS* (associated with homocystinuria (HOM)).^7^ It is unclear how well the patient cohort was characterized, as well as how many of the identified rare variants are known to be associated with disease or how strongly they are predicted to be pathogenic. The authors suggest the need for a prospective study to document the prevalence of such IEMs in patients with psychosis.

**Table 1.** Demographic characteristics of the study sample (n=2 046)

| **Characteristic** | **Schizophrenia** | **Bipolar Disorder** | **Major Depressive Disorder** | **Total** |
| --- | --- | --- | --- | --- |
| **Number of Samples** | 1 132 | 719 | 195 | 2046 |
| **Male : Female** | 663 : 469 | 336 : 383 | 107 : 88 | 1 106 : 940 |
| **%** | 59 : 41 | 47 : 53 | 55 : 45 | 54 : 46 |
| **Ethnicity, (%)** |  |  |  |  |
| *Caucasian* | 888 (78) | 611 (85) | 157 (81) | 1 656 (81) |
| *African* | 80 (7) | 37 (5) | 13 (7) | 130 (6) |
| *Eastern Asian* | 63 (6) | 17 (2) | 10 (5) | 90 (4) |
| *South Asian* | 61 (5) | 25 (4) | 6 (3) | 92 (5) |
| *Other* | 40 (4) | 29 (4) | 9 (5) | 78 (4) |

Beyond the diagnosis of patients with IEMs to allow for targeted treatments, there is emerging evidence that heterozygous carriers for autosomal recessive IEMs can present with milder or limited phenotypes. For example, there are case reports of carriers of single NPC mutations presenting with delirium and paranoid SCZ, and Parkinsonian tremors without biochemical or other systemic indications of classic NPC.^8, 9^ Bauer *et al.* (2013) observed a high frequency (4·8%) of heterozygous *NPC1* and *NPC2* gene variants, and identified three affected individuals, in their cohort of 250 patients with neurological and psychiatric symptoms.^10^ The authors suggest the possibility of a penetrant phenotype for heterozygous NPC carriers, but cite a lack of strong evidence to date to support this theory. Similar to NPC, there is some evidence that heterozygous *ATP7B* mutation carriers may also be at risk for neuropsychiatric phenotypes. Demily *et al.* (2017) found that 19% of 269 psychiatric patients studied had low ceruloplasmin and serum copper levels, and identified four heterozygous *ATP7B* mutation carriers, but no individuals confirmed to be affected with WD.^11^ Other case reports and series describe heterozygous WD carriers presenting with Parkinsonian tremors and psychiatric symptoms.^12, 13^ Significantly, the neurological and psychiatric symptoms of a heterozygous *ATP7B* mutation carrier with acquired hepatocerebral degeneration, and normal copper and ceruloplasmin levels, were almost completely ameliorated by treatment with penicillamine, a copper-chelating agent commonly used in the treatment of WD.^14^

The high heritability, heterogeneous symptomatology, and treatment resistance rates seen in SCZ, BPD, and MDD are suggestive of the involvement of intrinsic factors, such as underlying genetic disorders. Recognition of the clinical findings that may be associated with carrier status for recessive disorders and the potential for targeted treatment further increase the impact of identifying pathogenic IEM variants in psychiatric patients.

In this study, we test the hypothesis that psychiatric populations are enriched for pathogenic variants associated with treatable genetic disorders by performing next-generation sequencing (NGS) in large cohorts of well-defined SCZ, BPD, and MDD patients for pathogenic variants in genes associated with four treatable IEMs, NPC, WD, HOM, and acute intermittent porphyria (AIP).

## MATERIALS AND METHODS

### Samples

A total of 2046 DNA samples from SCZ (n=1132, 100 with tardive dyskinesia), BPD (n=719), and MDD (n=195) patients were included in this study. Sample characteristics were described previously elsewhere.^15, 16^ A subset of samples (n=1166) was obtained from the Individualized Medicine: Pharmacogenetic Assessment & Clinical Management (IMPACT) project at the Centre for Addiction and Mental Health (CAMH; Toronto, Canada), which consists of patients taking psychiatric medication who have experienced inadequate response, side effects, or may be switching to new psychiatric medication (http://impact.camhx.ca/en/home.php). The demographic characteristics of the sample sets are shown in Table 1. Research ethics board approval for this study was obtained through CAMH.

### DNA Sequencing

Primers for the five targeted genes (*NPC1, NPC2, ATP7B, HMBS, CBS*) were designed on the Ion AmpliSeq™ Designer (https://www.ampliseq.com/browse.action; supplementary Table 2). DNA samples were purified using Agencourt® AMPure® XP (Beckman Coulter Life Sciences, Indianapolis, IN, USA) and quality controlled for amplifiable DNA quantity using TaqMan® RNase P Detection Reagents Kit (ThermoFisher Scientific Inc., Waltham, MA, USA). NGS of the genes was carried out using a pooled matrix-based strategy (supplementary Figure 1) on the ION PROTON™ System for NGS (ThermoFisher Scientific Inc., Waltham, MA, USA). Known and predicted pathogenic variants identified by NGS were validated by Sanger sequencing (for primers, see supplementary Table 3).

**Table 2.** Known pathogenic variants (n=28) identified in the study cohort*

| Sample ID | Dx | Gene | Nucleotide Position | Protein Position | Exon | Variant Type |
| --- | --- | --- | --- | --- | --- | --- |
| 196 | SCZ | *NPC1* | c.1812dupT | --- | 12 | frameshift insertion |
| 483 | MDD | *NPC1* | c.3104C>T | A1035V | 21 | missense |
| 174 | SCZ | *NPC1* | c.3182T>C | I1061T | 21 | missense |
| 823 | BPD |  |  |  |  |  |
| 155 | SCZ | *NPC1* | c.3477+4A>G | --- | 22 | splice |
| 821 | MDD |  |  |  |  |  |
| 1648 | SCZ | *NPC1* | c.3560C>T | A1187V | 23 | missense |
| 729 | BPD | *NPC1* | c.3598A>G | S1200G | 24 | missense |
| 433 | SCZ | *NPC2* | c.88G>A | V30M | 2 | missense |
| 161 | SCZ |  |  |  |  |  |
| 1047 | BPD | *ATP7B* | c.406A>G | R136G | 2 | missense |
| 1603 | BPD | *ATP7B* | c.1915C>T | H639Y | 7 | missense |
| 843 | SCZ | *ATP7B* | c.2972C>T | T991M | 14 | missense |
| 4498 | BPD | *ATP7B* | c.2978C>T | T993M | 14 | missense |
| 538 | SCZ |  |  |  |  |  |
| 372 | BPD | *ATP7B* | c.3053C>T | A1018V | 14 | missense |
| 601 | SCZ | *ATP7B* | c.3069T>C | Thr1023= | 15 | missense |
| 847 | SCZ |  |  |  |  |  |
| 868 | MDD | *ATP7B* | c.3209C>G | P1070R | 15 | missense |
| 111 | SCZ | *ATP7B* | c.3688A>G | I1230V | 18 | missense |
| 14 | SCZ |  |  |  |  |  |
| 310 | SCZ | *ATP7B* | c.3955C>T | R1319* | 20 | nonsense |
| 659 | BPD | *ATP7B* | c.4092_4093delGT | --- | 21 | frameshift deletion |
| 3820 | BPD | *CBS* | c.1471C>T | R491C | 16 | missense |
| 291 | MDD | *HMBS* | c.176C>T | T59I | 4 | missense |
| 2742 | BPD | *HMBS* | c.569C>T | T190I | 9 | missense |
| 219 | SCZ | *HMBS* | c.962G>A | R321H | 14 | missense |
| 2346 | SCZ | *HMBS* | c.973C>T | R325* | 14 | nonsense |

**Table 3.** Predicted pathogenic variants (n=20) identified in the study cohort

| Sample ID | Dx | Gene | Nucleotide Position | Protein Position | Exon | Variant Type | Pathogenicity Score* | Polar Bond Disruptions | Affected Conserved Regions / Putative Effect | ACMG Classification |
| --- | --- | --- | --- | --- | --- | --- | --- | --- | --- | --- |
| 337<br>2365 | SCZ<br>SCZ | *NPC1* | c.180G>T | Q60H | 2 | missense | 3 | loss of 3·0 Å PB | N-terminal luminal loop/NPC1 domain (residues 55-165)<sup>38</sup> | likely pathogenic |
| 772 | SCZ | *NPC1* | c.873G>T | W291C | 6 | missense | 4 | --- | loss of hydrophobic side chain leading to disruption of protein folding | likely pathogenic |
| 318<br>871 | SCZ<br>SCZ | *NPC1* | c.2792A>T | N931I | 18 | missense | 4 | no change in PB | cysteine-rich luminal loop (residues 927 to 958)<sup>38</sup>, gain of hydrophobic side chain leading to disruption of protein folding | likely pathogenic |
| 503 | BPD | *NPC1* | c.3196A>G | T1066A | 21 | missense | 5 | loss of 2·1 Å PB | cysteine-rich luminal loop containing a ring-finger motif (residues 855-1098)<sup>38</sup> | likely pathogenic |
| 2812 | SCZ | *NPC1* | c.3556C>T | R1186C | 23 | missense | 5 | loss of 2·3 Å, 2·6 Å, and 3·1 Å PBs | loss of PBs between alpha-helices leading to instability between secondary structures<sup>39</sup> | likely pathogenic |
| 2279 | SCZ | *NPC1* | c.3811G>C | E1271Q | 25 | missense | 3 | --- | Near Di-leucine motif, which is necessary for lysosomal targeting | likely pathogenic |
| 810 | SCZ | *NPC2* | c.454dupT | X152delinsL | 5 | frameshift insertion | --- | --- | Loss of stop codon leading to non-stop decay of mRNA | pathogenic |
| 347<br>1505 | SCZ<br>BPD | *CBS* | c.1484C>T | T495M | 16 | missense | 4 | loss of 2·6 Å and 3·4 Å PBs | loss of PBs leading to disrupted protein folding and function | likely pathogenic |
| 76 | SCZ | *CBS* | c.1642C>T | R548W | 17 | missense | 4 | loss of 3·4 Å PB | loss of PBs leading to disrupted protein folding and function | likely pathogenic |
| 1833 | SCZ | *ATP7B* | c.372C>A | S124R | 2 | missense | 3 | --- | HMA domain 1 | likely pathogenic |
| 9<br>320 | BPD<br>BPD | *ATP7B* | c.442C>T | R148W | 2 | missense | 4 | --- | HMA domain 2 | likely pathogenic |
| 34 | SCZ | *ATP7B* | c.925A>G | M309V | 3 | missense | 3 | --- | HMA domain 3 | likely pathogenic |
| 955 | SCZ | *ATP7B* | c.1660A>G | M554V | 8 | missense | 3 | --- | HMA domain 5 | likely pathogenic |
| 670 | BPD | *ATP7B* | c.2737G>C | V913L | 15 | missense | 3 | --- | highly conserved amino acid position | likely pathogenic |
| 206 | SCZ | *ATP7B* | c.3337C>T | R1113W | 18 | missense | 4 | --- | loss of positively charged side chain leading to disruption of polar bond and secondary structure | likely pathogenic |
| 649 | SCZ | *HMBS* | c.71G>C | G24A | 2 | missense | 4 | no change in PB | loss of a compact side chain and gain of hydrophobic side chain leading to disruption of protein folding | likely pathogenic |

*Dx*, diagnosis; *SCZ*, schizophrenia; *MDD*, major depressive disorder; *BPD*, bipolar disorder; *NPC1*, Niemann pick C 1; *NPC2*, Niemann pick C 2; *ATP7B*, ATPase copper transporting beta; *CBS*, cystathionine-beta-synthase; *HMBS*, hydroxymethylbilane synthase; Å, Angstrom; PB, polar bond(s); ---, no protein structure was available to perform mutagenesis or mutation could not be modelled within the available protein structure; HMA, heavy metal-associated. \*'Pathogenicity Score' denotes the total number of software (SIFT, PolyPhen2, MutationTaster, MutationAssessor) which predicted the given variant to be damaging (SIFT score of $\leq 0.01$ , PolyPhen2 score of $\geq 0.80$ , MutationTaster score of 'disease causing' or 'disease causing automatic', MutationAssessor score of 'medium' or 'high', M-CAP score of $> 0.025$ ). All pathogenicity software used are only designed to predict pathogenicity of non-synonymous substitution variants.

### Bioinformatic Analysis

Bioinformatic analysis was performed on the CAMH Specialized Computing Cluster with in-house scripts (refer to supplementary documents for details). Pathogenicity of sequence variants was interpreted according to American College of Medical Genetics and Genomics (ACMG) guidelines.^17^

### Statistical Analysis

Statistical comparison of observed variant frequencies for NPC, WD, HOM, and AIP to expected frequencies based on general population carrier frequencies was performed using the exact binomial test, and comparison to the Genome Aggregation Database (gnomAD: http://gnomad.broadinstitute.org/) whole-exome sequenced population (WES; n=123 136) was performed using the Fisher’s exact test in R version 3·4·0.^18^ Exon coverage and depth plots are available on the gnomAD website. The gnomAD consortium intends to release a non-psychiatric subset in the near future; this was not available at the time of our analysis nor a release date mentioned. Statistical comparison of treatment response in subjects with and without pathogenic variants was performed using the Fisher’s exact test in R. All statistical test performed were 2-sided and all tests were Bonferroni corrected.

### Protein Modelling of Variant Substitutions

Protein modelling was performed using the PyMOL Molecular Graphics System, Version 1·8 (Schrödinger LLC, USA). Protein Data Bank (PDB) protein structures 3JD8, 4L3V, and 3EQ1 were used to model eight unique predicted pathogenic variants within NPC1, CBS, and HMBS proteins, respectively.^19-21^ The first, most likely, rotamer was chosen to depict the amino acid change. Protein modelling could not be performed for ATP7B and NPC2 variants due to a lack of complete protein structures in the PDB.

## RESULTS

### Variants Identified

Figure 1 summarizes the variants identified through bioinformatic analysis and validated by Sanger sequencing. Exon coverage and depth of all five genes sequenced averaged 400x per sample across all exons. A total of 1531 variants were identified in the five genes following filtration and annotation. Of these, 28 known pathogenic (22 missense, two missense leading to a stop gain, two splice, one insertion, and one deletion; Table 2 and supplementary Table 4), and 20 predicted pathogenic (one insertion leading to a stop loss, 19 missense; Table 3 and supplementary Table 5) variants were validated by Sanger sequencing. Notably, four known and one predicted pathogenic variants were found in the *HMBS* gene, strongly suggesting that the individuals carrying these variants are affected with AIP, an autosomal dominant disorder.

**Figure 1.**
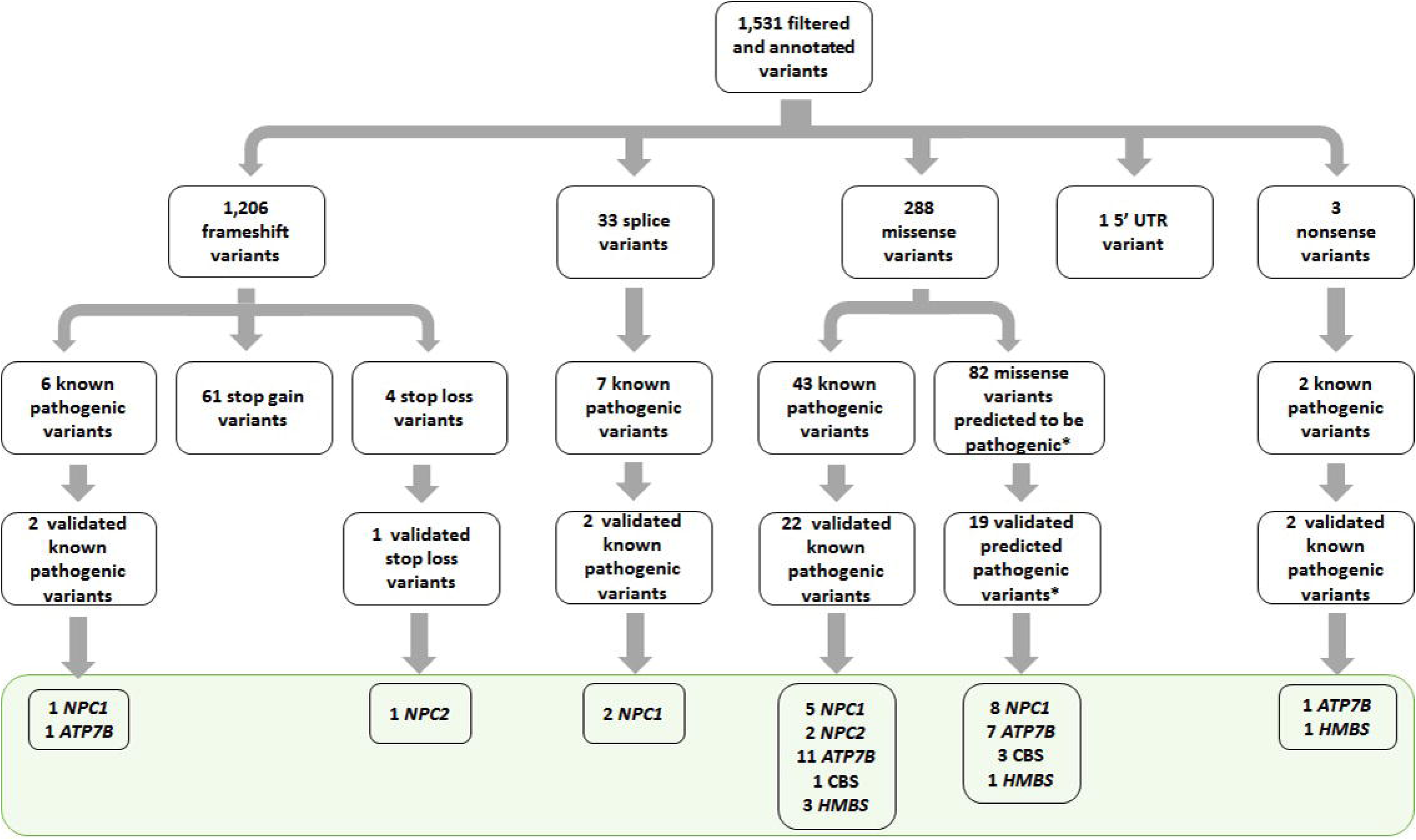
Diagram depicting the filtered and annotated variant breakdown of all 2046 SCZ, BPD and MDD samples. The grey box contains the final Sanger sequencing-validated predicted pathogenic variants identified. *NPC1*, Niemann pick C 1; *NPC2*, Niemann pick C 2; *ATP7B*, ATPase copper transporting beta; *CBS*, cystathionine-beta-synthase; *HMBS*, hydroxymethylbilane synthase. Variant types that did not undergo additional analysis are discussed in the limitations.

**Table 4.**
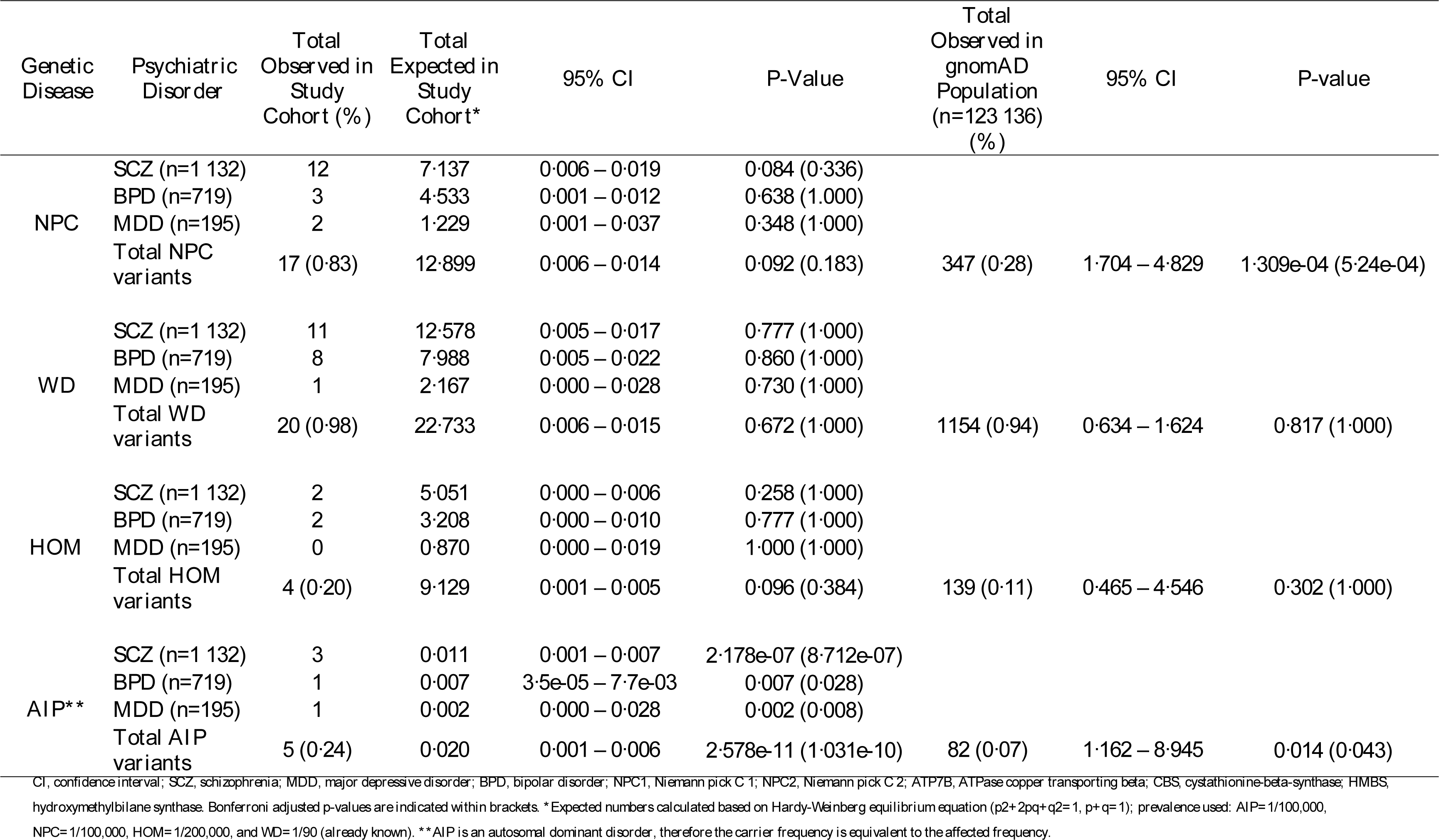
Prevalence of pathogenic variant frequencies for the selected IEMs in the study cohort compared to expected carrier frequencies in the general population and to gnomAD comparison population

### Prevalence of Pathogenic Variants in Psychiatric Populations

The carrier frequencies for the autosomal recessive disorders NPC, WD, and HOM, and the predicted rate of individuals affected with AIP in our SCZ, BPD, and MDD cohorts are shown in Table 4. Overall, NPC carrier rate was found to be marginally higher in the SCZ cohort (1·06%) compared to general population (95% CI, 0·006–0·019; p=0·336) and significantly higher than in the gnomAD population (95% CI, 1·704–4·829; p=5·24e-04) populations. Ethnicity-matched variant frequencies from the gnomAD WES cohort are shown in supplementary Table 5 for each of the predicted pathogenic variants; all variants were either absent or found at very low frequency. Additionally, two SCZ patients were found to have homozygous N931I likely pathogenic variant. The predicted affected rate of AIP was found to be significantly enriched across the entire cohort of SCZ, BPD, and MDD patients (0·24%) relative to the general (95% CI, 0·001–0·006; p=1·031e-10) and gnomAD (95% CI, 1·162–8·945; p=0·043) populations.

### Protein Modelling

The protein locations of the known and predicted pathogenic variants identified in this study are shown in supplementary Figure 2. Protein modelling results are shown in Table 3 for variants where analysis was possible. These predicted structural modifications to the NPC1, CBS, and HMBS proteins are shown in supplementary Figure 3.

## DISCUSSION

Our study used rigorous genetic and bioinformatic analysis, including Sanger sequencing validation of variants identified by NGS, stringent variant analysis parameters and protein modelling, to screen for pathogenic variants associated with selected treatable genetic diseases in well-defined psychiatric cohorts. In our study, we found a significant enrichment of pathogenic gene variants associated with NPC in SCZ, and AIP across the entire cohort of SCZ, BPD, and MDD patients compared to both the expected frequencies based on general population prevalence and the gnomAD comparison population. The variant frequencies seen in the gnomAD cohort were higher than expected in the general population, likely due to the inclusion of individuals from psychiatric exome studies. There remains a significant enrichment of NPC and AIP pathogenic variants in our pure psychiatric cohort compared to the heterogeneous gnomAD population, potentially hinting at the extent of the impact of these IEM variants in the pathogenesis of the psychiatric disorders. The frequencies of WD and HOM pathogenic variants were similar to those expected in the general population, though there was an enrichment of WD carriers in the psychiatric cohort compared to the gnomAD population. Protein modelling for predicted pathogenic variants identified in NPC1, CBS, and HMBS demonstrated an overall disruption of polar bonds, and the majority of NPC1 and ATP7B variants are located in functional or conserved domains, providing substantial support for the likelihood of these variants being damaging to protein structure and function.

One of the major limitations of this study was the inability to perform adequate analysis of insertion and deletion variants due to the intrinsic limitations of the ION PROTON™ sequencing platform. The error rate of the sequencer in homopolymer DNA regions resulted in high false positive call rates for small insertions/deletions resulting in frameshift mutations.^22^ Despite this, our study still demonstrated a significantly higher frequency of pathogenic variants in selected IEM genes in our psychiatric cohorts. This is highly suggestive that the prevalence of these rare disease variants in psychiatric populations is, in fact, much higher.

The NPC carrier rate identified in this study (0·83%), particularly in the SCZ cohort, was higher than that previously reported by Wassif *et al.* (2016), who estimated a population carrier rate of 0·659% based on their predicted incidence rates of 1/92 104 and 1/2 858 998 for NPC1 and NPC2, respectively.^23^ Their results were based on analysis of large exome data sets, with variants classified as pathogenic when predicted by at least two pathogenicity prediction algorithms used, while our analysis was based on more stringent criteria of pathogenic predictions by at least three algorithms, supporting the likelihood that the carrier rate is truly higher. Moreover, our NPC carrier frequency is likely to be an underestimation given the study limitations described. The estimated disease prevalence of NPC in our psychiatric cohort based on our detected carrier frequency is 0.002%, twice what is expected in the general population (0.001%).^24^ Two patients in the study cohort were found to carry the likely pathogenic N931I variant in the homozygous state, potentially representing affected individuals. Available medical history was insufficient to determine the likelihood of these individuals being affected with NPC and follow-up with biochemical testing will need to be pursued. Nevertheless, along with the carrier rate seen, the data support that NPC gene variants are enriched in psychiatric patients. Individuals with NPC exhibit cholesterol disturbances that can negatively affect the homeostasis of myelinated axons, which are lipid-rich and highly lipid-dependent brain structures, suggesting a possible mechanism by which *NPC1* and *NPC2* mutations could be contributing to mental illnesses.^25^ Controlled studies of heterozygous *NPC1(+/-)* mice show motor dysfunction, anxiety-like behaviour, and neurodegeneration, possibly through tau protein mechanisms, supporting the hypothesis that heterozygous NPC carrier states increase the risk for neuropsychiatric phenotypes.^26, 27^ Miglustat, the only approved treatment for NPC, has been shown to lower total tau levels in the cerebrospinal fluid of NPC patients, suggesting the possibility that treatment of heterozygous NPC carriers with miglustat could improve neuropsychiatric symptoms and prevent further neurodegeneration.^28^ Future studies as to the functional impact of being a heterozygous carrier for NPC variants will be of great interest in determining exact mechanisms for an association with psychiatric symptomatology. Biochemical assays in NPC carriers, such as by oxysterol or bile acid profiles, or filipin staining of fibroblasts, could be informative. Taken together, NPC should be considered more carefully in the psychiatric clinical setting, especially in patients with SCZ.

We did not detect any difference in carrier rate for WD in our cohort (0·98%) compared to the reported general population carrier rate (1·10%), or to the gnomAD population (0·94%).^29^ There is some suggestion that heterozygous carriers for WD may accumulate copper in the basal ganglia, and that associated neuropsychiatric symptoms are responsive to first-line treatments used in WD.^14, 30^ Given the potential for easily accessible and highly effective treatment, it is imperative that larger studies be undertaken to determine whether there is true enrichment of *ATP7B* pathogenic variants in psychiatric populations, particularly in patients with mood and affective disorders, which are more common in WD, as well as to further characterize the full spectrum of carrier phenotypes.

We did not detect any enrichment of HOM carriers within our psychiatric cohorts (0·20%) compared to an estimated carrier frequency of 0·45% using a conservative disease prevalence of 1 in 200 000 in the general population.^31^ There was also no enrichment compared to the gnomAD population (0·11%). The true carrier prevalence in the general population for HOM remains unknown. There are currently no known phenotypic associations with heterozygous *CBS* mutation carriers; however, no large-scale studies have been performed to investigate this possibility.

The most significant enrichment seen in our study cohort was for pathogenic variants associated with AIP, with a frequency of 0·24% across the entire cohort of SCZ, BPD, and MDD patients compared to the general population frequency of 0·001% (95% CI, 0·001–0·006; p=1·031e-10) and gnomAD population frequency of 0·07% (95% CI, 1·162–8·945; p=0·043).^32^ Other studies using biochemical screening have suggested a prevalence of up to 0.39% for AIP in psychiatric populations, though it is difficult to ascertain selection bias in some older studies.^33-35^ Cederlöf *et al.* (2015) reported a four-fold risk for manifesting AIP patients to develop SCZ or BPD, while their first-degree relatives had a two-fold risk; the authors suggest the possibility of psychiatric disorders as a latent presentation of AIP.^36^ Another study found a significant correlation of generalized anxiety with porphyrin metabolite levels in latent and manifest AIP patients.^37^ Together, our study and previous literature support that primary psychiatric presentations of AIP are more common than previously thought. Because AIP is an autosomal dominant disorder, carriers of pathogenic variants in *HMBS* are *de facto* affected with the disorder. Although AIP is an incompletely penetrant disorder, patients with the enzyme deficiency are at increased risk of acute porphyric crises, typically comprising abdominal pain, hypertension, neuropathy, and psychosis, at some point in their lifetime. Patients presenting initially with only psychiatric symptoms are therefore at risk for multi-systemic involvement. Recurrent attacks can lead to chronic health problems, including neurological, cardiovascular, and pain symptoms, which are entirely avoidable, as AIP is an eminently treatable disorder. Avoidance of lifestyle-related triggers, such as alcohol, smoking, and fasting, as well as porphyrinogenic medications, and the administration of glucose and hematin are highly effective in preventing and stemming porphyric attacks; these treatments have significantly reduced the morbidity and mortality associated with AIP.^38^ Molecular diagnosis of AIP in pre-symptomatic individuals allows for preventative management and reduces the likelihood of porphyric attacks to only 5%, making the timely genetic diagnosis of AIP all the more crucial.^38^

Pathogenic variants associated with genetic diseases could provide at least a partial explanation for treatment resistance to typical pharmacological therapies for psychiatric disorders. In individuals for whom drug response data was available (31/49), it is of interest to note that more than half (58%, 18/31) of patients, including 11 SCZ and seven BPD patients, identified in this study to have pathogenic variants in the disease genes were responding poorly to their psychiatric medications at the time of study recruitment. Of these poor responders, seven of 11 SCZ patients were taking last-line anti-psychotic treatment (i.e. clozapine, olanzapine), and six of seven BPD patients were non-responsive to lithium. In comparison, an analysis of treatment response in 725 patients without identified pathogenic variants for whom this information was available showed that only 16% (115/725) were poor treatment responders. This significant difference (58% versus 16%; 95% CI, 3.285–16.735; p=2.452e-07) illustrates a potential role of IEM variants in high treatment-resistance rates seen within psychiatric disorders.

While biochemical confirmation of the genetic diagnosis will need to be sought for these patients, particularly those suspected to be affected with AIP, identification of an underlying genetic disorder reveals targeted therapy that can quickly and effectively alleviate current and future symptomatology. In addition, there is emerging evidence to suggest that heterozygous carriers for recessive disorders, such as NPC and WD, may exhibit milder or limited phenotypes. Identification of a genetic diagnosis, or carrier status for a genetic disease, in an individual has important implications for family planning and recurrence risks for family members.

Our study results provide proof of principle that pathogenic genetic variants associated with treatable genetic diseases are enriched in primary psychiatric patient populations. Identification of such variants provides a plausible explanation for treatment non-responsiveness in a subset of psychiatric patients. The results of this study, much like mental illness and genetic diseases, have wide-reaching implications for patients seen across all specialties. This novel approach to identification of genetic variants involved in psychiatric pathogenesis leads to the potential for development of new, targeted therapies to treat mental health disorders. More in-depth studies with larger patient cohorts and rigorous genomic screening for variants associated with genetic diseases, particularly treatable ones, will be invaluable in determining the true prevalence of these disorders. Further study of the psychiatric phenotypes associated with carrier status for recessive disorders, or as latent manifestations of dominant or X-linked disorders, will also contribute to greater understanding of the pathogenesis of difficult-to-treat psychiatric disorders. Awareness of and ascertainment for these disorders will be critical in ensuring patients are accurately diagnosed and treated, with further-reaching implications for family planning and counselling of family members.

## ACKNOWLEDGEMENT

The authors would also like to acknowledge Dr. Kirti Mittal, Sajid Shaikh, Anna Mikhailov, and Natalie Freeman for their technical assistance during this study. Computations were performed on the CAMH Specialized Computing Cluster, which is funded by The Canada Foundation for Innovation, Research Hospital Fund. V.S. was funded by an Ontario Graduate Scholarship. J.S. was funded by the Brain and Behavior Foundation NARSAD Young Investigator Grant. The study was also funded by an Actelion Pharmaceuticals Canada Unrestricted Research Grant. None of the funding sources had any role in the study design, collection, analysis, and interpretation of data, writing of the report or the decision to submit the paper for publication.

