## Supplementary Materials for "Enrichment of gene variants associated with treatable genetic disorders in psychiatric populations"

**Supplementary Methods**

**Bioinformatic Analysis**

Bioinformatic analysis was performed on the CAMH Specialized Computing Cluster with in-house scripts. In detail, Ion Torrent fastq files were aligned using the TMAP aligner to the hg19 (GRCh37) reference genome after removing adapter sequences. Freebayes was used to call variants for the pooled targeted sequencing approach, using 0.005% threshold for the minimum number of reads in a pileup to be considered an alternate allele.^1^ Freebayes was also set to detect pooled inputs where variants in a single pool represented 32 samples. The variants from sequence pools and reshuffled sequence pools were then crossed to identify which variants occurred twice in the different pools sets. VCFtools was used to identify commonly occurring variants in different pool sets and only variants that were sequenced twice were kept for annotation. Variants were annotated using ANNOVAR using the hg19 reference and up-to-date annotations as of June 25, 2017 and avsnp147. After annotation, variants were filtered for the most likely disease-causing mutations, including frameshift and nonsense mutations in the coding regions of *NPC1*, *NPC2*, *ATP7B*, *CBS* and *HMBS*. Annotated variant lists were also searched for known pathogenic variants from ClinVar, Human Gene Mutation Database Professional 2017, disease-specific databases (<https://medgen.medizin.uni-tuebingen.de/NPC-db2/index.php>; <http://www.wilsondisease.med.ualberta.ca/>), and review of literature.^2^ Missense variants were analyzed for pathogenicity using SIFT, PolyPhen-2, MutationTaster, Mutation Assessor and MCAP. ^3-7^ If a variant was classified as ‘damaging’ or ‘likely damaging’ (SIFT score of ≤0.01, PolyPhen2 score of ≥0.80, MutationTaster score of ‘*disease causing*’ or ‘*disease causing automatic*’, MutationAssessor score of ‘*medium*’ or ‘*high*’, MCAP score of ≥0.025) by at least three of the pathogenicity prediction software, and had an allele frequency below 0.005 in the gnomAD whole-exome sequenced population (n = 123 136), that variant was further analyzed for amino acid conservation using the publically available ConSurf Server (<http://consurf.tau.ac.il/2016/>) using the default parameters and Bayesian method of conservation score calculations.^8^^,9^ Variants were classified according to the ACMG guidelines.^10^ Known and predicted pathogenic variants were validated by Sanger sequencing (The Centre for Applied Genomics, The Hospital for Sick Children, Toronto, Canada). A control analysis was run using the same analytical pipeline on the gnomAD (<http://gnomad.broadinstitute.org/>) whole-exome sequenced population.

**Statistical Analysis**

Exact binomial test and Fisher’s exact test were performed with the *binom.test* and *fisher.test* scripts, respectively.

**Supplementary Tables**

|  | **Niemann-Pick disease type C** | **Wilson disease** | **Homocystinuria** | **Acute intermittent porphyria** |
| --- | --- | --- | --- | --- |
| **Disease Prevalence** | 1/100 000^11^ | 1/30 000^12^ | 1/200 000^13^ | 1/100 000^14^ |
| **Carrier Frequency*** | 1/158·62 (0·63%) | 1/90 (1·11%) | 1/224·11 (0·45%) | --- |
| **Genes Implicated** | *NPC1; NPC2* | *ATP7B* | *CBS* | *HMBS* |
| **Genetic Inheritance** | AR | AR | AR | AD |
| **Molecular Pathway Affected** | Cholesterol transport | Copper transport | Methionine metabolism | Heme biosynthesis |
| **Molecular Consequence** | Cholesterol accumulation in lysosomes | Copper accumulation in liver and bloodstream | Homocysteine accumulation | Porphobilinogen accumulation |
| **Psychiatric Symptoms** | Dementia, psychosis, (manic-) depression | Psychosis, depression | Acute psychosis, OCD, personality disorders, affective/BPD symptoms | Hallucinations, paranoia, depression |
| **Non-Psychiatric Symptoms** | Hepatosplenomegaly, vertical supranuclear gaze palsy, ataxia, dystonia, cataplexy | Liver cirrhosis, parkinsonism | ID, skeletal abnormalities, thromboembolism, lens detachment | Abdominal pain, peripheral neuropathy, seizures |
| **Treatment** | Miglustat | Diet modifications, penicillamine, liver transplant | Vitamin B6/B12, folate and betaine supplements | Avoidance of triggers, glucose and hemin administration, liver transplant |

**Supplementary Table 1. Examples of genetic diseases that present with primary psychiatric symptoms**

*NPC1*, Niemann pick C 1; *NPC2*, Niemann pick C 2; *ATP7B*, ATPase copper transporting beta; *CBS*, cystathionine-beta-synthase; *HMBS,* hydroxymethylbilane synthase; AR, autosomal recessive; AD, autosomal dominant; OCD, obsessive compulsive disorder; ID, intellectual disability. **Carrier frequency was calculated based on Hardy-Weinberg equilibrium equation (p2+2pq+q2=1, p+q=1); prevalence used: AIP=1/100,000, NPC=1/100,000, HOM=1/200,000, and WD=1/90 (already known).*

**Supplementary Table 2. Amplicon primers for next generation sequencing**

| **Gene** | **Amplicon Forward Primer** | **Amplicon Reverse Primer** | **Chr** | **Start Position*** | **Stop Position*** |
| --- | --- | --- | --- | --- | --- |
| *NPC1* | GTCAGGAAGGAAGAAGGCGTC | CCAGACTCCATAAGTCCCGC | chr18 | 21166462 | 21166633 |
| *NPC1* | CTGGCTTCTTAGAAGGCATGTGA | GAGTCTAGAATCTAGAGGAAGCAGCTA | chr18 | 21120960 | 21121234 |
| *NPC1* | CTCCGCTGCTTCTGAAGTACAA | CAGAGTGTTCACACTCTCTCCTATTC | chr18 | 21119736 | 21120008 |
| *NPC1* | GCCAGTTCCTTGGCTTTAAAACA | CAGCAAGCATCTTGTCTCCTTTTTC | chr18 | 21141262 | 21141536 |
| *NPC1* | AAACAAAGAATAAATGGAAAGCTGAGCATT | GAATGTGTCTTAGTTCACTGAGGAATGT | chr18 | 21151954 | 21152226 |
| *NPC1* | CAGGTAGCCAGCTCCTTCTTTC | TTAACACAAGGCAGCAAGAAATGG | chr18 | 21123349 | 21123610 |
| *NPC1* | ACTCTTCAGTCACTGAGGAGGAT | GGTAATTAGCACCCATCCTCAGAAC | chr18 | 21114339 | 21114611 |
| *NPC1* | GTGACCACAGATGGAACAAGC | CCTCCTGCTGCTACTGTGTC | chr18 | 21166006 | 21166278 |
| *NPC1* | GTTCAACATCATTCACTTTCTTGAAACCT | ACTGACAAACACATTTACCAGCCATA | chr18 | 21136026 | 21136290 |
| *NPC1* | AAGTTGGAATGAAGAAAATAGATGTAGGCA | ACAGTGATGTCTTCACCGTTGTAATTAG | chr18 | 21124749 | 21125018 |
| *NPC1* | CAAGGATACAGCGTTCAGACTGA | AGTGAGAGCGAGCTTTAATGAGG | chr18 | 21115448 | 21115706 |
| *NPC1* | AAACTTCACAGGGCAAGGTCTT | CGACTTCTTTGTGTATGCCGATTAC | chr18 | 21134594 | 21134772 |
| *NPC1* | CTAGGTCTATTTCTAGCTCAATGTAAGACG | AAAGCAACATGTTCTTCACAGTGTTC | chr18 | 21111342 | 21111616 |
| *NPC1* | AAAGTGTATCTACAACCTCAACTGTCAC | GACTTGGGAAGCAGTATTACTAGATCTG | chr18 | 21111753 | 21112000 |
| *NPC1* | GCTTTACCTGTAAGGAAATACTCGGT | CTCTCTTGACACCCAGGATTCTTTC | chr18 | 21116630 | 21116870 |
| *NPC1* | GCAGTGTTAAACAGAACTTTGATGGT | TTGGACGCCATGTATGTCATCA | chr18 | 21140010 | 21140283 |
| *NPC1* | CCCTTAGACACAGTTCAGTCAGGAT | CTGCAGAAATAAGAAAAAGTCTCTCTCTCT | chr18 | 21112127 | 21112283 |
| *NPC1* | CTGACGAACACGCAGTAATGAAG | AGATAGCAACTAATGCTTTCCCTGTTC | chr18 | 21136434 | 21136665 |
| *NPC1* | AACATGTGGACCTTGTTCAGCT | GAGTGGACAATATCACTGACCAGTTC | chr18 | 21119096 | 21119357 |
| *NPC1* | GCAAGTGTCTAGCTTCCCACAA | GGTATGTGTCTAATTTTCTGCATGCTT | chr18 | 21127911 | 21128136 |
| *NPC1* | GTCTTAGCCCAGTCCTCTCCTA | AAAAATCTGGAGACCTATTCTTCTAACAGT | chr18 | 21118424 | 21118687 |
| *NPC1* | GAGGTAAGAAATTAACAAAACTGCCCAA | CCTGATGTCTTGAGGCCCTTCTA | chr18 | 21131472 | 21131745 |
| *NPC1* | GAAAGATTTGGTAAAGGAGAAGGTACCT | CATGTAGTTTTTCTTTTGACTGTTAGCAGT | chr18 | 21119293 | 21119563 |
| *NPC1* | GAATAATTACAGAGGATCTTGTGATCAGCA | GCACTTCTTGTTGAAATTTACCATTGAGAC | chr18 | 21153351 | 21153604 |
| *NPC1* | TGCTTGAAACACCTACGTGCAT | AATCCTGCTTTTTGTGTGTGCTTAAG | chr18 | 21120323 | 21120594 |
| *NPC1* | CCACTGAGGAAACGAATGCTCT | GGGATTACAGGAATGTCCCAAAAACA | chr18 | 21137007 | 21137232 |
| *NPC1* | CTGTCCTGATGCCAGCTGTAAA | GGCCCTATTATGTGTGAGATCATGC | chr18 | 21148739 | 21149012 |
| *NPC1* | CGCTAGCTGCTTCCTCTAGATT | TGTACATGCACATGAACATAAGACCTG | chr18 | 21121205 | 21121479 |
| *NPC1* | CCCAAGGCTAGGGAAATATATAGAAACA | GAGAAGTTTCTTACTTAGCTGTCAGTTAGT | chr18 | 21124952 | 21125178 |
| *NPC1* | CCCTGAAACTTGAACAGATGCTGA | AACCCTGTAACTAATTGGTGATTGTGT | chr18 | 21124259 | 21124533 |
| *NPC1* | CCTGCGCTGGACACAGTAG | CGACGACGCCTTCTTCCT | chr18 | 21166250 | 21166486 |
| *NPC1* | AGGATAGAATTCCCTTTCAGTAATGTCCT | TTACAGGTTGGTAAAAGTGGTTTCTAACA | chr18 | 21113254 | 21113528 |
| *NPC1* | GGAGGTCCAAAGGGTACATCAG | GAAACCCTGGCTGTGTCATTTTC | chr18 | 21136232 | 21136490 |
| *NPC1* | CAAGCGCCAGACTTGGTATCTTA | CTTGGTCAACATGTTTGGAGTTATGTG | chr18 | 21115317 | 21115508 |
| *NPC1* | CACTTACCGTACGCAGTACAGA | ACGTGTTTCTGGGTTTGCTTATTTTTAAAA | chr18 | 21134715 | 21134989 |
| *NPC1* | CTCAGGCCTTCACAGAGACTTTA | TCACCGAAACCATGGGCATTAA | chr18 | 21116437 | 21116688 |
| *NPC1* | GCCACATCTAACTGGCAATTAAATCTCTT | GAGTAAGCCATCCCACAAGTTCTATA | chr18 | 21111551 | 21111817 |
| *NPC1* | GCTGGGAGAAGTTTAGTGTCCT | CGAACGGCTTCTAAATTTCTAGCC | chr18 | 21111940 | 21112187 |
| *NPC1* | AACACAAGCAAAAACGCCATGTA | ATTGAGATTTGTACTCAACACAATTCCTTT | chr18 | 21140228 | 21140480 |
| *NPC2* | CTTATGGCACTGATTTAGTTTCAGTCTGA | GAAGTTTGCTACTGTACATCTAGGATTCAT | chr14 | 74947320 | 74947582 |
| *NPC2* | CCATTCCCATGCTTATTCCAACAC | CAGAGCACCTTCCCATTAGGTG | chr14 | 74952952 | 74953226 |
| *NPC2* | GTAGCTGCCAGGAAACGCAT | CAGCTGTGGTTACTGGTGACA | chr14 | 74959958 | 74960101 |
| *NPC2* | TGAAAAAGTCATGTCTTCAGTGCACT | TGGTTGTCTCATGTCTCTTTTTCTGT | chr14 | 74946533 | 74946806 |
| *NPC2* | CCTCAGAACTCTAATCCAGTCCCAA | AGGTTATTTTCTTTGCCATCTGATTCTCT | chr14 | 74951059 | 74951332 |
| *NPC2* | GCTAACCAAGTGCTGCATTTAATGA | CCACTGAGCTGGGACATTACATC | chr14 | 74946745 | 74947019 |
| *NPC2* | CCAGCCCATTCCAGTTAGGTAG | TTCTTTCCCGAGCTTGGAACTT | chr14 | 74959751 | 74960010 |
| *NPC2* | GCGGTCACAAGACAAACCTGT | AGAGGTGCTCTAAAGGGAAGGAA | chr14 | 74960024 | 74960276 |
| *ATP7B* | GAGCTATAAGACACAAAGAGAAAAGGAGA | CAGTACCACTCTGATTGCCATTG | chr13 | 52548008 | 52548282 |
| *ATP7B* | TTGCCTGATATCTGCAGAAAACTGT | TTGCATGGTTTTTAGTTCACAGTGAAATT | chr13 | 52515152 | 52515426 |
| *ATP7B* | AGGTCTATTGCAATGTCAATACAACATG | CTCTAACACCACGCTTGTGACT | chr13 | 52531556 | 52531830 |
| *ATP7B* | GACATGGTGAGGAATAAAAGAGCATTG | TGGTCAGTGAGTTGTGGTTGTT | chr13 | 52518183 | 52518457 |
| *ATP7B* | AACTGTCTGATTTCCCAGAACTCTTC | GCTGAGCAAGTGACAGTTGTCT | chr13 | 52524063 | 52524334 |
| *ATP7B* | CCCTACTGTTAAAATCCTATCCTCCTCT | CCTTTTAGATGGTCAAAGTGTAAGGAGTTT | chr13 | 52506677 | 52506911 |
| *ATP7B* | CGGAGGCATAAGTGATGCCATT | CTTTCACAGGCTTTCCTTGATCCT | chr13 | 52539088 | 52539227 |
| *ATP7B* | GGCTGGTACAAGAAGGGTCATA | AAAGCACTAACCCAAAGAGACCTTTA | chr13 | 52548434 | 52548673 |
| *ATP7B* | GCTGAGTGAGACTTTGACTCTCA | ACTGTGAAATATGTGCCATCGGTT | chr13 | 52548819 | 52549061 |
| *ATP7B* | GCTGATGATGCCTTTCAAATTGGA | CTGGGATGTTGTAGAAAATATTTGGTTTCA | chr13 | 52549092 | 52549366 |
| *ATP7B* | CTAGCTTTTTAGAAAGGACCAGAGTGA | CCTGGTCCTGGCACTGATTTAT | chr13 | 52511188 | 52511461 |
| *ATP7B* | GGGCAATGAACACAAAGAGCATG | TGAACTCTCCTCCCTACTTGCT | chr13 | 52532476 | 52532730 |
| *ATP7B* | TGATGGACGTCTGGAAAGCAAA | TAGAGTTCTGGGAGCTTCCTTATTGA | chr13 | 52520551 | 52520676 |
| *ATP7B* | CCACCTGTCATCCATGCCTATG | TCTAGAATGGCTCAGATGCTGTTG | chr13 | 52509060 | 52509205 |
| *ATP7B* | TAAGTTCAACATGGGCGTTCATCT | CTAACCCTCTTGGAAACCACAGT | chr13 | 52544599 | 52544873 |
| *ATP7B* | CTTCTCTGGCTGTGATCTGTCTC | CAACTTTGAATCATCCGTGTGAAGAG | chr13 | 52585437 | 52585691 |
| *ATP7B* | AATGATCAGCCTAGTCAGAAAACAACA | AACTCAGAATGCACTTGATTCAGGA | chr13 | 52507014 | 52507254 |
| *ATP7B* | CTTGTCCATTGGCTATGTCATCCT | GGGAGCAAACTAGTAGAGTTGGATTAAA | chr13 | 52507392 | 52507638 |
| *ATP7B* | GGAGGCTGTGTTTTCCTCCTAT | CCTGTGGTGCTTGAAACGTTTG | chr13 | 52507794 | 52507989 |
| *ATP7B* | TCTATTGAGAAGCCAACACTCCATG | CCAATGTCCTTGTGGTCTTTGCT | chr13 | 52508108 | 52508299 |
| *ATP7B* | CACAGGAGAGAAAAGGAACAGACTATG | CTCTGCTTGGAGTATTTAGGATGACT | chr13 | 52508353 | 52508591 |
| *ATP7B* | GCCAAGGACTAGAGTCCAAGACA | GGAGCAGTACATCTGATGACTTCAG | chr13 | 52508634 | 52508907 |
| *ATP7B* | CCCAGGTAGAGGAAGGGACTTAG | AATTTCCAAAGCTGAAAAGTGCTTTCT | chr13 | 52535914 | 52536141 |
| *ATP7B* | CTTCGGACAGTCCTCTTGGAAA | AGAATAAAGGGAAGAAAGTCGCCAT | chr13 | 52511474 | 52511748 |
| *ATP7B* | CATGTGACCTGACAGCTGCTAT | TAACAGCTGGCCTAGAACCTGA | chr13 | 52524348 | 52524608 |
| *ATP7B* | TTGTCTCTAACTGCTTTTATGAGCTTTACA | CATTGCAAGTGTGGTATCTTGGTG | chr13 | 52513131 | 52513402 |
| *ATP7B* | TTTTTGGTCCTGATGAAACTGTTCTC | CCCTGGATATGTCCAGTCATCCT | chr13 | 52508529 | 52508778 |
| *ATP7B* | TCTGTGGTTTGACCCACCTCTA | CCCTCTTGGCTTACAGTTTCCT | chr13 | 52516464 | 52516736 |
| *ATP7B* | GCCCAGTGAATCTAAGATATGAAAGAACA | CTTCATAGGTTGTAATTTCCCATGGTCT | chr13 | 52523707 | 52523979 |
| *ATP7B* | GCAGCATTTGTCCCAGGTGAAT | CTAGGTGTGAGTGCGAGTTCTT | chr13 | 52509610 | 52509863 |
| *ATP7B* | TATCTGAGGGCCACACACAGCAT | GCAGCATCTGATATATCTGTGTTGCT | chr13 | 52534226 | 52534499 |
| *ATP7B* | TTACTAATCACAAAGATGGATGTGTCCAAA | GAGTGTTACAGCCATGACCTGA | chr13 | 52542500 | 52542774 |
| *ATP7B* | GGCTCTCAGGCTTTTCTCTCAA | ACATCTCCCAGACAGAGGTGAT | chr13 | 52520333 | 52520601 |
| *ATP7B* | CCTTCAATGGAATGGACACAGGAT | CAAGTGTCCTTGGAGAACAAAACTG | chr13 | 52548220 | 52548491 |
| *ATP7B* | CCCAAGGTCTCAGAATTATTAAAATTCTGG | TTGAAGGCAAGGTCCGGAAACT | chr13 | 52548607 | 52548874 |
| *ATP7B* | CCCAATTTGATGGCAAACCTGT | CCAGTCATGTGTGAAGTCCATTG | chr13 | 52549005 | 52549149 |
| *ATP7B* | CTGGGATTTCAGAAGTAGTGACCA | GAGGAGCCCTGTGACATTCTTC | chr13 | 52532257 | 52532531 |
| *ATP7B* | CCTGAAGTCATCAGATGTACTGCT | GCCTGACCTGGAGAGGTATGAG | chr13 | 52508882 | 52509156 |
| *ATP7B* | GAGCTTATTTCCATGGGAAAAGTTGAAG | TGTGTCCACAACATAGAGTCCAAAC | chr13 | 52538871 | 52539145 |
| *ATP7B* | GTGTACCATCTGTAGTTTGCACCAT | CCTGAAACCTCTTGTTCTGAAAAACATATT | chr13 | 52544814 | 52544958 |
| *ATP7B* | ACACTCCAGAGCATTGGAGAAG | TGCTTTCTTCCTGCATAGTCTGTTC | chr13 | 52508241 | 52508415 |
| *ATP7B* | ACATCAGTTGACGGCACACTTT | CCAGATCAGAGAAGAATTCGGTGT | chr13 | 52585238 | 52585509 |
| *ATP7B* | GGCAGATTTTTAGAGGAATGACAGGA | ATTGTGTCCTCTCTTTATGCTTGCT | chr13 | 52506799 | 52507072 |
| *ATP7B* | AGTGAAACTAACCATCCAAGGTGAAG | TTACAGGCAAGGAAACAGGCTCCAA | chr13 | 52507193 | 52507459 |
| *ATP7B* | AACTATTTTGTGTGGGAGAAAAGGGT | CTTGTGTGGCTTGGAGGAAATG | chr13 | 52507574 | 52507848 |
| *ATP7B* | CCTGCACACATACGTTTCCCAT | TCTTCTTCAAGTTGAGGAGAGTTCTTTTT | chr13 | 52507942 | 52508180 |
| *ATP7B* | CCTACCTGCTGCAATGGGTATC | GACGTCGTCCTTATCAGAGTGAG | chr13 | 52511407 | 52511629 |
| *CBS* | CCAGTCTACTTTGTCTCGACCTT | GGGATCGGCTACGACTTCATC | chr21 | 44482892 | 44483104 |
| *CBS* | ACCGTGAGGAATGACAGCTTTC | GGGCTGAGTGTGTTTTCAATGATT | chr21 | 44483930 | 44484183 |
| *CBS* | GCCACTCATTAACCAGCGAGTT | TCCTGAATAATTGTGGACTCCTCTGT | chr21 | 44488483 | 44488756 |
| *CBS* | GTGACTGCGCATCTGTTTGAG | GGCAGAGGACTTCCATGTGTG | chr21 | 44478212 | 44478483 |
| *CBS* | CCCGAATGCTGGTCAAAGGA | GCGGATGATTGAGGATGCTGA | chr21 | 44486165 | 44486432 |
| *CBS* | TGTCACTGCGAGTGTGCAT | CGCTGCGTGGTCATTCTG | chr21 | 44480328 | 44480591 |
| *CBS* | ATAAGGACAAACGCTCTCGCA | GAAGGGCTTTCTGAAGGAGGAG | chr21 | 44479111 | 44479380 |
| *CBS* | CGCAGTGACACTCCTCAGAAC | GGTGGTGGACAAGTGGTTCAA | chr21 | 44482287 | 44482506 |
| *CBS* | GCTCCTTGGCTTCCTTATCCTC | AGTTCTTCCTGGGCTTCTCTGA | chr21 | 44492183 | 44492434 |
| *CBS* | GCCTCACCTGGTCTAGGATGT | CCAGAGAAGATGAGCTCCGAGAA | chr21 | 44485490 | 44485749 |
| *CBS* | GTTTTACTTGGTTAACTTCTTGCCCTT | GCGCAAGGCGACTGTTCT | chr21 | 44496351 | 44496621 |
| *CBS* | CTTCTCTCTTTTGCCTTTAATCCACTCT | CTTCGCTTTCCTGAGCCCTAAA | chr21 | 44473662 | 44473936 |
| *CBS* | GGCATAAAGACTGGGTGTCACT | ACATCCTGGAGATGGACCACTT | chr21 | 44476760 | 44476963 |
| *CBS* | TGACATGCCTGAAAATACCATGCA | GAAAGTGAACAATCAGCGGCATTT | chr21 | 44473247 | 44473513 |
| *CBS* | CCGGGACCCAGTTGAGATC | TGCTCTGCCACGAGACATTG | chr21 | 44495816 | 44496054 |
| *CBS* | CCACTCACACTGGATCTGCT | TCTCACTCCACAGAAAACTCGTG | chr21 | 44476905 | 44477061 |
| *CBS* | ACTCCTGCCCTCCAGGTTAT | CCCAGCATGCCTTCTGAGA | chr21 | 44492040 | 44492309 |
| *CBS* | GGGCTCTGGACTCGACCTA | GATCATTAACAGGCAGTTGTTAACGG | chr21 | 44483043 | 44483303 |
| *CBS* | GTGGAGCTGGGCAGACAGAAC | TGCGAGAGCGTTTGTCCT | chr21 | 44478888 | 44479131 |
| *CBS* | GCAGCCAGGGATAAATGCAAT | GCGGCTGAAGAACGAAATCC | chr21 | 44485270 | 44485539 |
| *CBS* | GATGTCGGCTCGATAATCGTGT | GGCAATTTTTCAGAACCCACAGA | chr21 | 44486364 | 44486620 |
| *CBS* | AAGCCGTGTCTTACATGTAGTTCC | GTGCACAATTCATGCATACGTGT | chr21 | 44480537 | 44480730 |
| *CBS* | CGCTTCACCCTCCTTTGATTCC | ACATAACCATTGTTGACATTAACCAAAGTC | chr21 | 44493313 | 44493535 |
| *CBS* | GAGTCGAACCTGGCATTGGT | GGCTATCGCTGCATCATCGT | chr21 | 44485567 | 44485773 |
| *CBS* | CGGTCTTACCAGGGCTTCTTC | ACCAGTGAGGTCCAGGAGAG | chr21 | 44479327 | 44479519 |
| *CBS* | CAAAGGTGAACGCCTCCTCAT | GTTGGAACTGGAAAGTCTGCAGA | chr21 | 44482457 | 44482671 |
| *CBS* | CATTCCCACGCCCTGTTGA | GTTTCAAGCTCATCAGTAAAGGTTCC | chr21 | 44496205 | 44496414 |
| *CBS* | GAGGTGGTGCCTACACAACTTT | CAGGTAGGATGAACACAGGCAA | chr21 | 44473457 | 44473720 |
| *CBS* | AGGCCAGGCAGTTACCAATC | AACCCACTGCCTCGTTCTC | chr21 | 44473886 | 44474119 |
| *HMBS* | AGATTCTTGATACTGCACTCTCTAAGGT | TTTAAGCCCAGCAGCCTATCTG | chr11 | 118959816 | 118960090 |
| *HMBS* | GTCAGCCAGCTAGAGAGGGAAAG | GGATGACTGTAAGGCAGAAAGGA | chr11 | 118962027 | 118962299 |
| *HMBS* | GGAAATTCCAGTCCCTTCAGGAT | GACGGGCTTTAGCTATAGGCAA | chr11 | 118959245 | 118959515 |
| *HMBS* | TCAAGAAATACCAGTGAGTTGGCAA | AGATTTTAACACTAGGCAGTCACTGTTC | chr11 | 118960605 | 118960876 |
| *HMBS* | AGGAGGACTGTGGCATTTCTTC | CCATCTTCATGCTGTATGAGGGAA | chr11 | 118963556 | 118963830 |
| *HMBS* | CCATAGAAGCTGCACTACTTGCT | GCTTGGAAAGTAGGCTGTGTGT | chr11 | 118955463 | 118955737 |
| *HMBS* | GAAAGATCAGGCCTGATGTCCT | GAGTTAGCACTGTATACAGAGCATTCA | chr11 | 118963074 | 118963335 |
| *HMBS* | GGAGCCAAAAACATCCTGGATGT | GCTTGGACTTCTCTAAAGAGATGAAGC | chr11 | 118963943 | 118964208 |
| *HMBS* | ACTGACAACTGCCTTGGTCAAG | AATCACTGGCTTGGAAGAAAGGAA | chr11 | 118958659 | 118958893 |
| *HMBS* | GCCAGACTCACACTTAGGCCTA | CCCTCCCTGAGAATGCTATTCTG | chr11 | 118962715 | 118962989 |
| *HMBS* | GGAAAGGAACAGTGACTGCCTA | CCTCTAGACCTTGTCTTTTTCCTTGG | chr11 | 118960843 | 118961054 |
| *HMBS* | CATTCTTGTTGAATGTTGTGTATGGATAGG | GGGACTACCTAGAAACCTGGGAT | chr11 | 118963335 | 118963606 |
| *HMBS* | GCAGGAACCAGGGATTATGTGC | TCCTTGGTAAACAGGCTTTTCTCTC | chr11 | 118959681 | 118959955 |
| *HMBS* | TTCCTTTCTTCCAAGCCAGTGAT | CTTCATACTAGGAACTAACCCTCTGAGT | chr11 | 118958870 | 118959118 |
| *HMBS* | ACAAGAGTGCATATAATCTCTTGTTCTCA | CAAACCAGTTAATGGGCATCGTTAA | chr11 | 118963763 | 118964001 |
| *HMBS* | GTTCAAGCCTTCCAGGGATTTG | GGCTTTGTGTTTGTTCCTATCTTCC | chr11 | 118964112 | 118964309 |
| *HMBS* | GGAGACCAGGAGTCAGACTGTA | CAATAGACGACTGAGGATGGCAA | chr11 | 118955610 | 118955867 |
| *HMBS* | CTTGAGAAGGTGTGCTTCCTGA | GGGAAAGGCAAAGGTTCACATGA | chr11 | 118960265 | 118960534 |

Chr, chromosome; *NPC1*, Niemann pick C 1; *NPC2*, Niemann pick C 2; *ATP7B*, ATPase copper transporting beta; *CBS*, cystathionine-beta-synthase; *HMBS,* hydroxymethylbilane synthase. **start and stop positions are based on genomic build hg19/ GRCh37.*

| **Gene** | **Exon** | **Forward Primer** | **Reverse Primer** |
| --- | --- | --- | --- |
| ***NPC1*** | 2 | CCA CCC TGC AAT AAC ATT TAA GG | GAA ATT TAC CAT TGA GAC CCT GG |
|  | 6 | GAA TAG CTG TAG GAC ACA ATA ATC | GTA CTC AAC ACA ATT CCT TTC TG |
|  | 12 | GGA ATA AGA ATA AAG AGG CAA AAA TAT G | GTA TCG TGA AAG TTA GGG AGA AG |
|  | 18 | CAA GAC AAG GTG GTA CTG AC | CTC TCT CCT ATT CTT TTA TCT TTC |
|  | 21 | GCC CTT TGC TGG GTA AAC C | GCT GAT ATT TTG CAA GAC CTG G |
|  | 22 | CAT CTT TAG GGT TTA CAT GGA ATC | GCA GTG GTG ACA GGA TGA AC |
|  | 23 | CTT TGT GGT GCG ACT CTG C | GAG CCA TCC TAA AGG AAG TG |
|  | 24 | GCC ACC CTT TTA AGA TGA GAA C | GGT TTC TAA CAC AGT ATC TCT TC |
|  | 25 | GTA AAC CGA CCG ACC CTT AG | CTA GCT CCC TTT CTC CTG C |
| ***NPC2*** | 2 | CAT TCC CAT GCT TAT TCC AAC AC | GTG GGC AGC CTA GCT GG |
|  | 5 | GAG CAG GAG AAG ACC ACA G | CTT GCC CTA GGG TTA TTG CC |
| ***ATP7B*** | 2 | CTT GCC TTC AAT GGA GCT GAC | GGA ACA AGG CAG TGC CAC TG |
|  | 3 | CAG GGC TCA CCT ATA CCA C | CAG GTC TTC CAG TTC TCA TTC |
|  | 7 | GGG TTC ACA TTA CAA GGG TAA AG | GTA AAG AAG TTG TAA GCA GAA AAC C |
|  | 8 | CCA CAC ACA GCA TGG AAG G | CTT AAA CTG TGT CCT CAG AAG G |
|  | 14 | GTT ATA CTT GAC TTC CTA TTC TAT G | CAA GTT CGT CAC GTT GTG TC |
|  | 15 | GAC CAC ACA GAG AAG GCT C | GAG ATT GAA CGA CAG AGG ATC |
|  | 18 | CTC ACG TGC AAC ACT ACA TGG | GTA TCT TGG TGC GGG GTG C |
|  | 20 | TCT AGC CAG CCA GTG AGT G | GAT GGG GTC AAT GAC TCC C |
|  | 21 | GCA TGC ACA CCA GGC TCC | CTC TCC CCA GAC CTA GGT G |
| ***CBS*** | 16 | CAT AAA GAC TGG GTG TCA CTG | CTT CTT TCC CAT CTC ACA CAC |
|  | 17 | CTC ATA GGC CGT AAA CAG GG | CCC CTC AGA CCA CAG CAC |
| ***HMBS*** | 2 | CTT TCT TCC AAG CCA GTG ATT C | GCT GTG AGC ATC ATA ACT GTT C |
|  | 4 | GGG CTG CTC CCA GTT CTG | CAC CAC ACT CTC CTA TCT TTA C |
|  | 9 | GTC CTT AGC AAC TCT CCA CAG | CCT ACG GTG TTA GAG GTG GG |
|  | 14 | GCT CAG ATA GCA TAC AAG AGA C | GTT GCT TGG ACT TCT CTA AAG AG |

**Supplementary Table 3. Primers used for Sanger sequencing**

*NPC1*, Niemann pick C 1; *NPC2*, Niemann pick C 2; *ATP7B*, ATPase copper transporting beta; *CBS*, cystathionine-beta-synthase; *HMBS,* hydroxymethylbilane synthase.

| **Sample ID** | **Dx** | **Gene** | **Nucleotide Position** | **Protein Position** | **Literature Reference** |
| --- | --- | --- | --- | --- | --- |
| 196 | SCZ | *NPC1* | c.1812dupT | --- | Jahnova *et al.* (2014)^15^ |
| 483 | MDD | *NPC1* | c.3104C>T | A1035V | Ribeiro *et al.* (2001)^16^; Pedroso *et al.* (2012)^17^ |
| 174 | SCZ | *NPC1* | c.3182T>C | I1061T | Yamamoto *et al.* (1999)^18^; Battisti *et al.* (2003)^19^; Efthymiou *et al.* (2015)^20^ |
| 823 | BPD |  |  |  |  |
| 155 | SCZ | *NPC1* | c.3477+4A>G | --- | Synofzik *et al.* (2015)^2^^1^ |
| 821 | MDD |  |  |  |  |
| 1648 | SCZ | *NPC1* | c.3560C>T | A1187V | Fancello *et al.* (2009)^22^ |
| 729 | BPD | *NPC1* | c.3598A>G | S1200G | Bauer *et al.* (2013)^23^; Wassif *et al.* (2016)^24^ |
| 433 | SCZ | *NPC2* | c.88G>A | V30M | Park *et al.* (2003)^25^; Alazami *et al.* (2015)^26^; Bell *et al.* (2011)^27^ |
| 161 | SCZ |  |  |  |  |
| 1047 | BPD | *ATP7B* | c.406A>G | R136G | [Mukherjee *et al.* (2014)](http://www.ncbi.nlm.nih.gov/sites/entrez?cmd=Retrieve&db=PubMed&list_uids=24094725&dopt=Abstract)^28^; Coffey *et al.* (2013)^29^ |
| 1603 | BPD | *ATP7B* | c.1915C>T | H639Y | [Gromadzka *et al.* (2005)](http://www.ncbi.nlm.nih.gov/sites/entrez?cmd=Retrieve&db=PubMed&list_uids=16283883&dopt=Abstract)^30^; [Braiterman *et al.* (2014)](http://www.ncbi.nlm.nih.gov/sites/entrez?cmd=Retrieve&db=PubMed&list_uids=24706876&dopt=Abstract)^31^ |
| 843 | SCZ | *ATP7B* | c.2972C>T | T991M | [Cox *et al.* (2005)](http://www.ncbi.nlm.nih.gov/sites/entrez?cmd=Retrieve&db=PubMed&list_uids=16088907&dopt=Abstract)^32^; [Drury *et al.* (2015)](http://www.ncbi.nlm.nih.gov/sites/entrez?cmd=Retrieve&db=PubMed&list_uids=26275891&dopt=Abstract)^33^; [Luoma *et al.* (2010)](http://www.ncbi.nlm.nih.gov/sites/entrez?cmd=Retrieve&db=PubMed&list_uids=20333758&dopt=Abstract)^34^ |
| 4498 | BPD | *ATP7B* | c.2978C>T | T993M | Lepori *et al.* (2007)^35^; Schushan *et al.* (2012)^36^ |
| 538 | SCZ |  |  |  |  |
| 372 | BPD | *ATP7B* | c.3053C>T | A1018V | [Loudianos *et al.* (1998)](http://www.ncbi.nlm.nih.gov/sites/entrez?cmd=Retrieve&db=PubMed&list_uids=9671269&dopt=Abstract)^37^; [Schushan *et al.* (2012)](http://www.ncbi.nlm.nih.gov/sites/entrez?cmd=Retrieve&db=PubMed&list_uids=22692182&dopt=Abstract)^36^ |
| 601 | SCZ | *ATP7B* | c.3069T>C | Thr1023= | [Genetic Services Laboratory, University of Chicago](https://www.ncbi.nlm.nih.gov/clinvar/submitters/1238/), SCV000246745.1 (ClinVar); [PreventionGenetics](https://www.ncbi.nlm.nih.gov/clinvar/submitters/239772/), SCV000301714 (ClinVar) |
| 847 | SCZ |  |  |  |  |
| 868 | MDD | *ATP7B* | c.3209C>G | P1070R | [Dong *et al.* (2016)](http://www.ncbi.nlm.nih.gov/sites/entrez?cmd=Retrieve&db=PubMed&list_uids=27022412&dopt=Abstract)^38^ |
| 111 | SCZ | *ATP7B* | c.3688A>G | I1230V | Davies *et al.* (2008)^39^; Denoyer *et al.* (2013)^40^; [Schushan *et al.* (2012)](http://www.ncbi.nlm.nih.gov/sites/entrez?cmd=Retrieve&db=PubMed&list_uids=22692182&dopt=Abstract)^36^ |
| 14 | SCZ |  |  |  |  |
| 310 | SCZ | *ATP7B* | c.3955C>T | R1319* | Thomas *et al.* (1995)^41^; Prella *et al.* (2001)^42^; Xiong *et al.* (2015)^43^ |
| 659 | BPD | *ATP7B* | c.4092_4093delGT | --- | [Shah *et al.* (1997)](http://www.ncbi.nlm.nih.gov/sites/entrez?cmd=Retrieve&db=PubMed&list_uids=9311736&dopt=Abstract)^44^ |
| 3820 | BPD | *CBS* | c.1471C>T | R491C | Kraus *et al.* (1999)^45^ |
| 291 | MDD | *HMBS* | c.176C>T | T59I | Schneider-Yin *et al.* (2008)^46^; Chen *et al.* (2016)^47^; Xiong *et al.* (2015)^4^^3^ |
| 2742 | BPD | *HMBS* | c.569C>T | T190I | Schuurmans *et al.* (2001)^48^ |
| 219 | SCZ | *HMBS* | c.962G>A | R321H | Schuurmans *et al.* (2001)^48^; Amendola *et al.* (2015)^49^; Chen *et al.* (2016)^47^ |
| 2346 | SCZ | *HMBS* | c.973C>T | R325* | Petersen *et al.* (1996)^50^ |

**Supplementary Table 4. Literature reference list for known pathogenic variants identified in the study cohort**

Dx, diagnosis; SCZ, schizophrenia; MDD, major depressive disorder; BPD, bipolar disorder; *NPC1*, Niemann pick C 1; *NPC2*, Niemann pick C 2; *ATP7B*, ATPase copper transporting beta; *CBS*, cystathionine-beta-synthase; *HMBS,* hydroxymethylbilane synthase.
**Supplementary Table 5. Variant frequencies in comparison population and conservation analysis of predicted pathogenic variants identified in the study cohort**

| **Sample ID** | **Dx** | **Gene** | **Nucleotide Position** | **Protein Position** | **Exon** | **Variant Type** | **Total gnomAD Exome Variant Frequency** | **Ethnicity Matched gnomAD Exome Variant frequency** | **Amino Acid Conservation Score*** |
| --- | --- | --- | --- | --- | --- | --- | --- | --- | --- |
| **Known Pathogenic Variants** | | | | | | | | | |
| 196 | SCZ | *NPC1* | c.1812dupT | --- | 12 | frameshift insertion | absent | absent | --- |
| 483 | MDD | *NPC1* | c.3104C>T | A1035V | 21 | Missense | 0.00000812 | 0.00000000 | --- |
| 174 | SCZ | *NPC1* | c.3182T>C | I1061T | 21 | Missense | 0.00021120 | 0.00039390 | --- |
| 823 | BPD |  |  |  |  |  | 0.00021120 | 0.00039390 | --- |
| 155 | SCZ | *NPC1* | c.3477+4A>G | --- | 22 | Splice | 0.00066310 | 0.00901500 | --- |
| 821 | MDD |  |  |  |  |  | 0.00066310 | 0.00008296 | --- |
| 1648 | SCZ | *NPC1* | c.3560C>T | A1187V | 23 | Missense | 0.00011390 | 0.00000000 | --- |
| 729 | BPD | *NPC1* | c.3598A>G | S1200G | 24 | Missense | 0.00084890 | 0.01184000 | --- |
| 433 | SCZ | *NPC2* | c.88G>A | V30M | 2 | Missense | 0.00220300 | 0.00113800 | --- |
| 161 | SCZ |  |  |  |  |  | 0.00220300 | 0.00113800 | --- |
| 1047 | BPD | *ATP7B* | c.406A>G | R136G | 2 | Missense | 0.00031300 | 0.00000000 | --- |
| 1603 | BPD | *ATP7B* | c.1915C>T | H639Y | 7 | Missense | 0.00005279 | 0.00010740 | --- |
| 843 | SCZ | *ATP7B* | c.2972C>T | T991M | 14 | Missense | 0.00121800 | 0.00052560 | --- |
| 4498 | BPD | *ATP7B* | c.2978C>T | T993M | 14 | Missense | 0.00007750 | 0.00013490 | --- |
| 538 | SCZ |  |  |  |  |  | 0.00007750 | 0.00013490 | --- |
| 372 | BPD | *ATP7B* | c.3053C>T | A1018V | 14 | Missense | 0.00005450 | 0.00004568 | --- |
| 601 | SCZ | *ATP7B* | c.3069T>C | Thr1023= | 15 | missense | 0.00013540 | 0.00000000 | --- |
| 847 | SCZ |  |  |  |  |  | 0.00013540 | 0.00029090 | --- |
| 868 | MDD | *ATP7B* | c.3209C>G | P1070R | 15 | missense | 0.00000813 | 0.00000896 | --- |
| 111 | SCZ | *ATP7B* | c.3688A>G | I1230V | 18 | missense | 0.00030460 | 0.00006497 | --- |
| 14 | SCZ |  |  |  |  |  | 0.00030460 | 0.00057290 | --- |
| 310 | SCZ | *ATP7B* | c.3955C>T | R1319* | 20 | nonsense | 0.00008122 | 0.00012530 | --- |
| 659 | BPD | *ATP7B* | c.4092_4093delGT | --- | 21 | frameshift deletion | absent | absent | --- |
| 3820 | BPD | *CBS* | c.1471C>T | R491C | 16 | missense | absent | absent | --- |
| 291 | MDD | *HMBS* | c.176C>T | T59I | 4 | missense | 0.00010960 | 0.00002685 | --- |
| 2742 | BPD | *HMBS* | c.569C>T | T190I | 9 | missense | 0.00004027 | 0.00008741 | --- |
| 219 | SCZ | *HMBS* | c.962G>A | R321H | 14 | missense | 0.00117400 | 0.00208600 | --- |
| 2346 | SCZ | *HMBS* | c.973C>T | R325* | 14 | nonsense | absent | absent | --- |
| **Predicted Pathogenic Variants** | | | | | | | | | |
| 337 | SCZ | *NPC1* | c.180G>T | Q60H | 2 | missense | 0.00028430 | 0.00043880 | 0.344 |
| 2365 | SCZ |  |  |  |  |  |  |  |  |
| 772 | SCZ | *NPC1* | c.873G>T | W291C | 6 | missense | 0.00007338 | 0.00112000 | 2.276 |
| 318 | SCZ | *NPC1* | c.2792A>T | N931I | 18 | missense | absent | absent | 0.314 |
| 871 | SCZ |  |  |  |  |  |  |  |  |
| 503 | BPD | *NPC1* | c.3196A>G | T1066A | 21 | missense | 0.00002437 | 0.00005372 | -0.978 |
| 2812 | SCZ | *NPC1* | c.3556C>T | R1186C | 23 | missense | 0.00001627 | 0.00002688 | -1.068 |
| 2279 | SCZ | *NPC1* | c.3811G>C | E1271Q | 25 | missense | 0.00001624 | 0.00000000 | -0.476 |
| 810 | SCZ | *NPC2* | c.454dupT | X152delinsL | 5 | frameshift insertion | absent | absent | --- |
| 347 | SCZ | *CBS* | c.1484C>T | T495M | 16 | missense | 0.00007177 | 0.00149100 | 1.539 |
| 1505 | BPD |  |  |  |  |  |  |  |  |
| 76 | SCZ | *CBS* | c.1642C>T | R548W | 17 | missense | 0.00011450 | 0.00006331 | 0.324 |
| 1833 | SCZ | *ATP7B* | c.372C>A | S124R | 2 | missense | 0.00001219 | 0.00002689 | 0.381 |
| 9 | BPD | *ATP7B* | c.442C>T | R148W | 2 | missense | 0.00117100 | 0.00899900 | 0.130 |
| 320 | BPD |  |  |  |  |  |  |  |  |
| 34 | SCZ | *ATP7B* | c.925A>G | M309V | 3 | missense | 0.00000406 | 0.00000896 | 2.269 |
| 955 | SCZ | *ATP7B* | c.1660A>G | M554V | 8 | missense | 0.00069540 | 0.00041300 | 0.414 |
| 670 |  | *ATP7B* | c.2737G>C | V913L | 15 | missense | absent | absent | -0.979 |
| 206 | SCZ | *ATP7B* | c.3337C>T | R1113W | 18 | missense | 0.00002437 | 0.00000895 | 0.135 |
| 649 | SCZ | *HMBS* | c.71G>C | G24A | 2 | missense | absent | absent | -0.964 |

*NPC1*, Niemann pick C 1; *NPC2*, Niemann pick C 2; *ATP7B*, ATPase copper transporting beta; *CBS*, cystathionine-beta-synthase; *HMBS,* hydroxymethylbilane synthase. *Conservation score is called using The ConSurf Server; the score depicts the relative conservation rate of each amino acid position within its gen;, the lowest score depicts the most conserved position in the given protein.

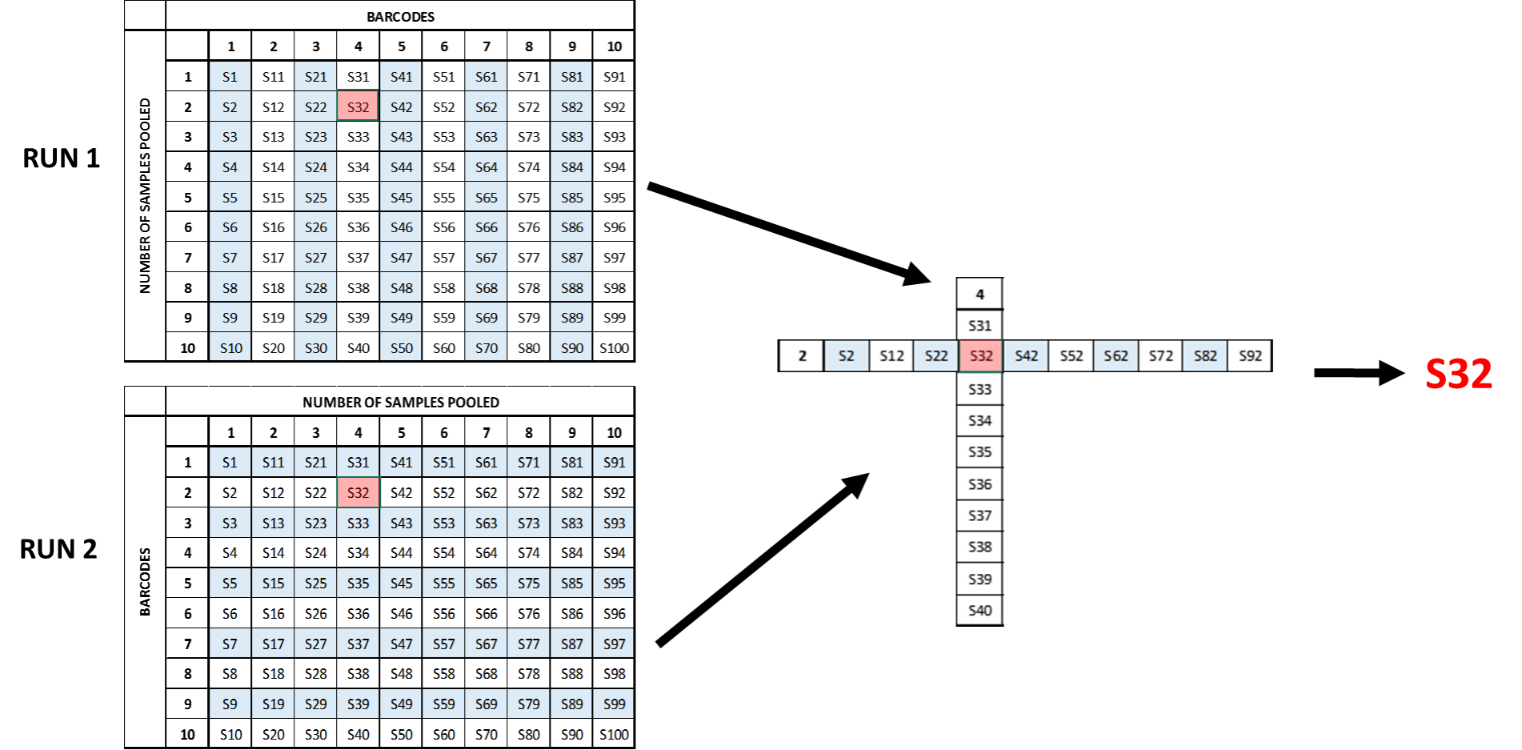
**Supplementary Figures**

**Supplementary Figure 1. Pooled matrix targeted sequencing**

To increase throughput of samples and cost-effectiveness, multiple samples are pooled under one barcode (e.g. samples S1 through S10 under barcode 1) and multiple barcodes are sequenced on one chip. In this example (Run 1), 100 samples are sequenced on a single chip by pooling 10 samples per barcode, with 10 barcodes in total. To determine the exact individual within a pool to have a specific variant, a second chip (Run 2) with the same samples is run with the sample pools shuffled, such that no two samples from the previous pool are re-pooled together in the second run. Through deduction from the barcodes identified in each run to contain the variant of interest, the specific sample carrying the particular variant can be identified. In the study, 32 samples were pooled together under one identifying barcode and prepared into one library according to the manufacturer’s protocol (Ion Ampliseq™ Library Preparation, Thermo Fisher ScientificInc., USA). A total of 1023 samples, along with a positive control (DNA from an NPC patient; GM18436, Coriell Institute for Medical Research Biorepository), in 32 libraries were sequenced concurrently. A total of four sequencing runs were performed to sequence 2046 samples (and positive control for each run).

**Supplementary** **Figure 2. Diagram depicting all known and predicted pathogenic variants**

R325*

R321H

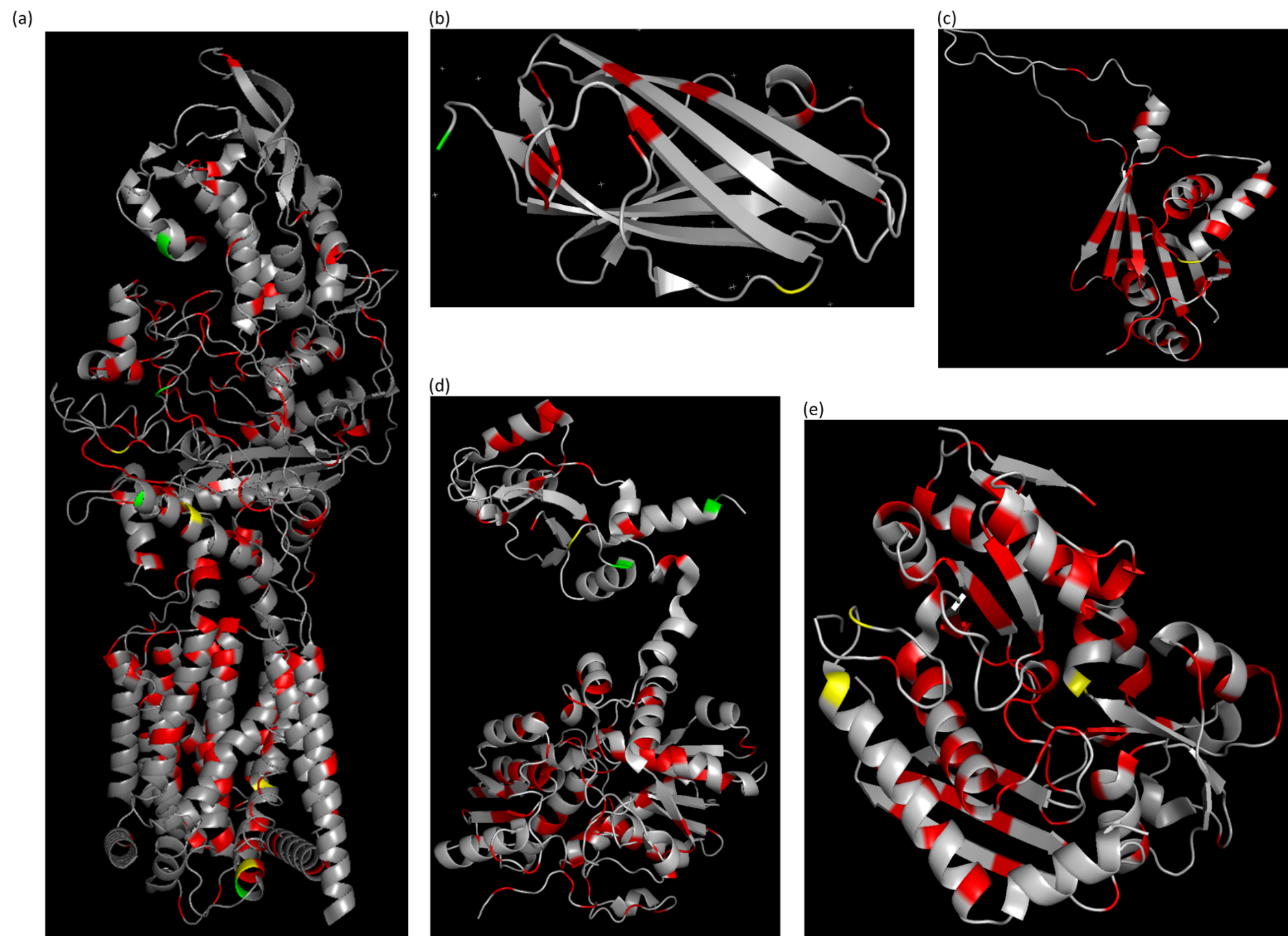

Q60H

N931I

A1035V

T1066A

I1061T

R1186C

A1187V

S1200G

V30M

X152delinsL

P1070R

R548W

R491C

T495M

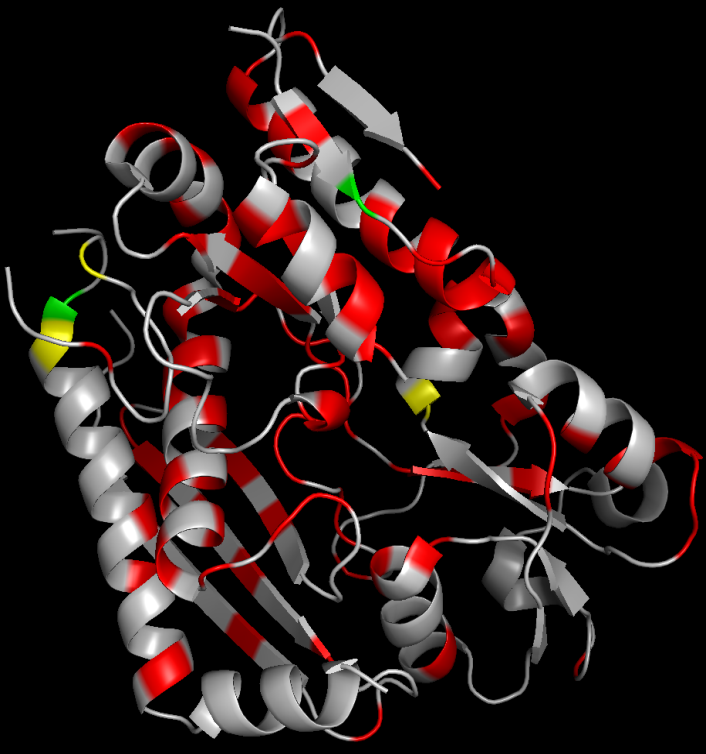

T190I

R321H

R325*

P324Q

G24A

All known pathogenic variants that were detected in this study are highlighted in yellow, predicted pathogenic variants detected in this study highlighted in green, and additional known missense mutations from HGMD in red. (a) NPC1; (b) NPC2; (c) N-domain of ATP7B; (d) CBS; (e) HMBS. Two NPC1 and 15 ATP7B variants could not be modelled due to incomplete crystal structure available.

**Supplementary Figure 3. Protein modelling of predicted pathogenic missense variants**

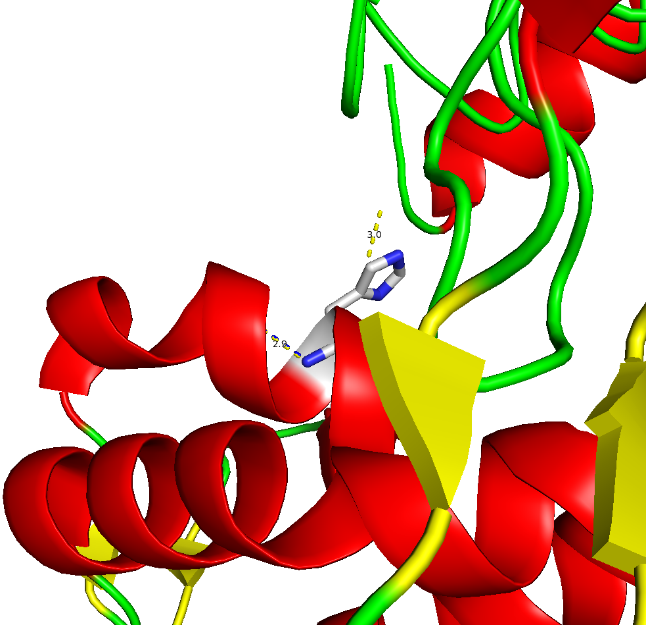

(a)

Loss of 3.0 Å polar bond

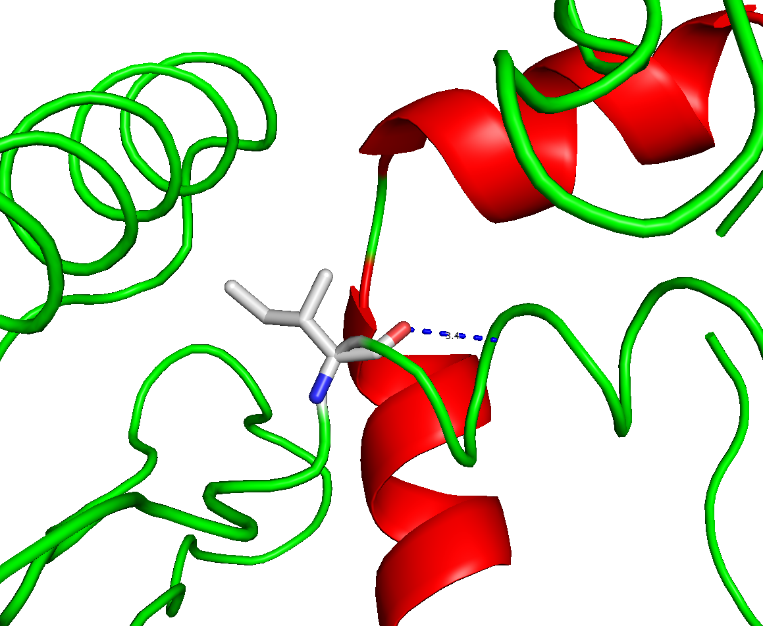

(b)

No change in

3.4 Å polar bond

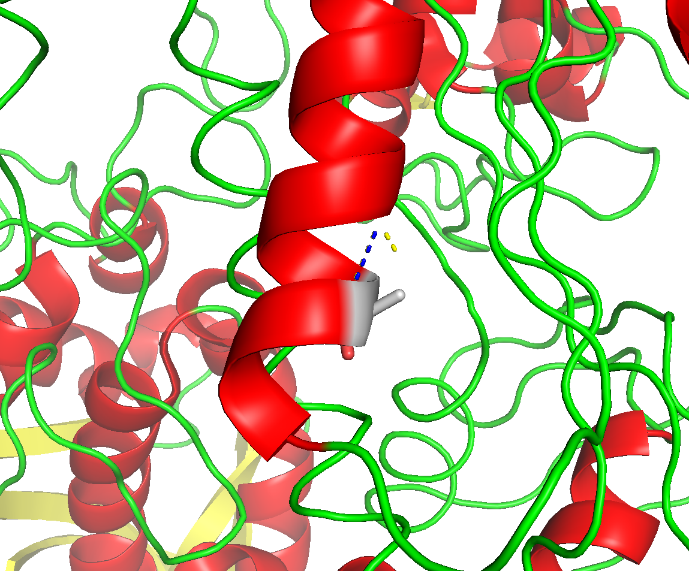

(c)

Loss of 2.1 Å

polar bond

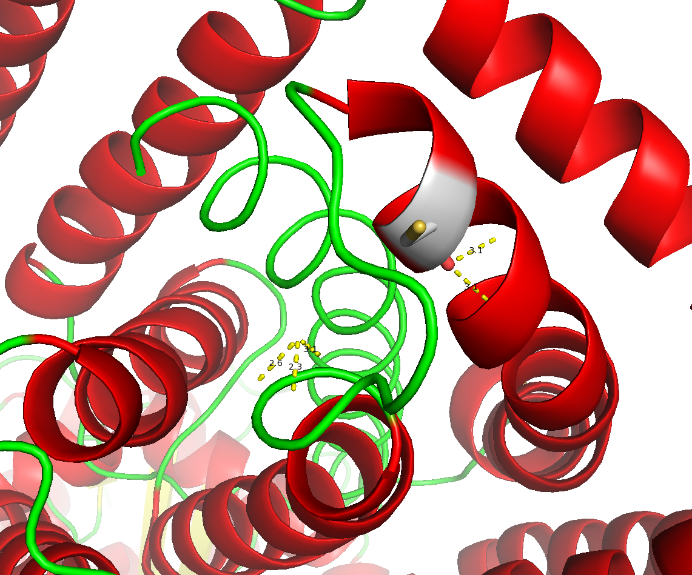

(d)

Loss of 2.3 Å, 2.6 Å, and 3.1 Å polar bonds

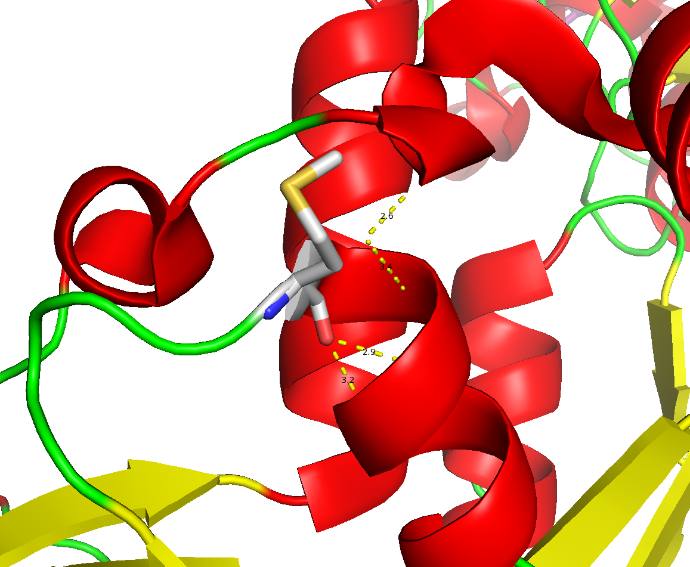

(e)

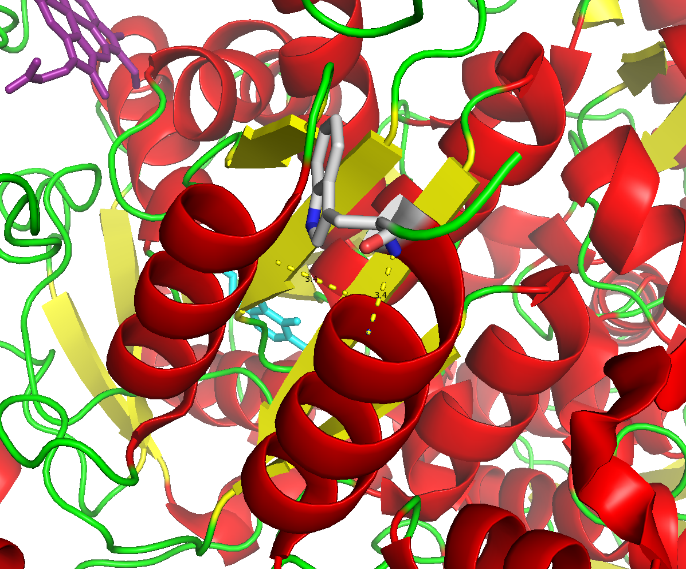

(f)

Loss of 3.4 Å polar bond

Loss of 2.6 Å and 3.4 Å polar bonds

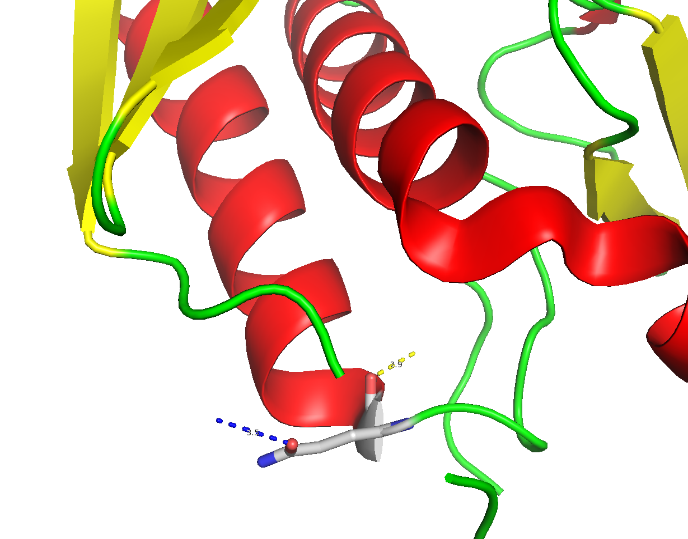

(f)

Loss of 3.9 Å polar bond

Gain of 3.5 Å polar bond

Protein modelling of predicted pathogenic missense variants depicting the polar bonds formed pre- (yellow dashed lines) and post- (blue dashed lines) substitution at the amino acid position of interest. (a) NPC1 Q60H; (b) NPC1 N931I; (c) NPC1 T1066A; (d) NPC1 R1186C; (e) CBS T495M; (f) CBS R548W. The NPC1 W291C and E1271Q variants could not be modelled with PyMOL, as these mutations were not captured in the crystalline protein structures available in PDB (Figure 2b). W291C and E1271Q are situated at the cytoplasmic side of the topological domain, and these substitutions could result in the loss of polar bonds necessary for proper formation of secondary structures and protein folding. As well, one predicted pathogenic variant was identified in NPC2, X152delinsL, which results in the loss of a stop codon. Without an in-frame stop codon, stop loss mutations can result in non-stop decay of transcript, resulting in the loss of a functioning protein.^51^ The seven ATP7B predicted pathogenic variants could not be modelled due to lack of a complete crystalline protein structure of ATP7B. Nevertheless, variants S124R, R148W, M309V, and M554V are all situated within heavy metal-associated (HMA) domains, each of which binds and transports one copper ion.^52^ HMA domains are highly conserved and play an important role in ATP7B protein functioning, supporting the disruptiveness of variants in these regions. Though the V913L variant was not located in a HMA domain, a pathogenic variant at the same amino acid position (V913I) has previously been reported.^53^ Similarly, though the HMBS G24A variant did not result in any change in polar bonds or fall within any known conserved regions, two AIP mutations have previously been identified at the same amino acid position (G24S and G24D).^54,^^55^
